# Increased sensitivity to myopia and altered retinal ON/OFF balance in a mouse model lacking *Dusp4*

**DOI:** 10.1101/2025.09.29.678242

**Authors:** Baptiste Wilmet, Christelle Michiels, Jingyi Zhang, Awen Louboutin, Sanja Boranijasevic, Helen Frederiksen, Jacques Callebert, Juliette Varin, The MyoTreat Consortium, Marie-Laure Gimenez, Said El Shamieh, Linda Horan, Robin Plevin, Serge Picaud, Catherine Morgans, Robert Duvoisin, Olivier Marre, Isabelle Audo, Christina Zeitz

## Abstract

Myopia, influenced by environmental and genetic factors, occurs when the emmetropization process fails to stop, causing excessive eyeball growth. Highly myopic animal models lacking a functional ON-pathway identified *Dusp4* as a potential gene implicated in myopia. Here, we used a mouse model lacking DUSP4 to gain a better understanding of its retinal role and the mechanisms implicated in myopia development. *Dusp4^-/-^* mice have a reduced basal level of retinal dopamine and a higher susceptibility to lens-induced myopia. *Dusp4* is expressed in ON-bipolar cells and a subset of OFF-bipolar cells in a light dependent manner. The absence of DUSP4 causes a hyperactivation of the MAPK/ERK pathway. *Dusp4^-/-^* mice showed reduced optomotor responses, increased ON-bipolar cell depolarization, reduced oscillatory potentials together with altered OFF and ON-OFF RGC responses to light flashes. These data provide insights into retina-driven mechanisms of myopization, nuancing the impact of ON and OFF pathways upon emmetropization.

## Introduction

Myopia is the most common ocular dysfunction worldwide, especially in Southeast Asia [1, 2]. It is characterized by an abnormal increase in the axial length during refractive development (i.e. emmetropization), leading to blurry distance vision but normal near vision. Previous studies revealed that both environmental factors such as light and genetic factors can be involved in myopia initiation and progression [3]. High myopia (<-6.00D) can lead to significant ocular complications that may cause severe vision loss and even blindness [4]. Given the major role of the retinal ON-pathway in perceiving light increments and the importance of light perception for proper emmetropization, studying myopia in the context of ON-pathway defects may offer new insight into the development of myopia.

In this respect, one form of inherited retinal disorders (IRDs), complete congenital stationary night blindness (cCSNB), caught our attention since it is associated with impairment of dim light vision and high myopia [5–7]. It is characterized by a complete loss of the b-wave upon electroretinography, in relation with ON-bipolar cell (ON-BC) pathway dysfunction [5]. ON-Cone Bipolar Cells (ON-CBCs) form direct excitatory synapses with dopamine-releasing amacrine cells (DACs) while rod-bipolar cells (RBCs) have indirect excitatory output to DACs through gap junctions to ON-CBCs. The retinal dopaminergic system, plays a critical role in proper emmetropization [8, 9]. Indeed, retinal levels of dopamine (DA) and its metabolite 3,4-Dihydroxyphenylacetic acid (DOPAC) are considered as hallmarks of myopia as they are reduced in a wide variety of experimental models of myopia [8–14]. The precise mechanisms through which dopaminergic system impact emmetropization remains under debate. The main endpoint of the signaling cascade implies remodeling of the scleral extracellular matrix, allowing the growth of the eyeball [15].

The role of ON, OFF pathways and ON/OFF balance upon emmetropization and susceptibility to myopia is currently under debate. To date, mouse models of pure ON pathway defect (*Nyx^-/-^*[16], *Gpr179^-/-^* [12], *Grm6^-/-^* [17] and *Lrit3^-/-^* [13]) all showed higher sensitivity to myopia, most probably due to the loss of excitatory input of DACs from ON-BCs, causing an altered retinal dopaminergic activity. In contrast, mouse models with a selective OFF pathway defect (*Vsx1^-/-^* [18]) were found not to be more susceptible to develop myopia nor reduced DA metabolism despite having a slightly altered contrast sensitivity [19]. In humans, hyperactivation of OFF pathway tends to cause choroidal thinning [20], a frequent compound of myopia. Recent research also suggested that myopia occurring in patients with pathological variants in *GJD2* primarily imply an altered OFF pathway [21]. In addition, mice depleted for *Aplp2* showed reduced b-wave and oscillatory potentials (OPs, thought to be mediated by ACs), higher sensitivity to myopia but also degeneration of OFF cone bipolar cells (OFF-CBCs) [22, 23]. ON and OFF pathways show a tight interplay as ON activity can inhibit OFF activity. All these findings led us to consider the effect of ON/OFF imbalance upon ocular elongation rather than exclusive alteration of ON or OFF pathways.

Our previous study performing whole RNA sequencing (RNAseq) on adult retinas from three cCSNB mouse models, *Gpr179^-/-^, Lrit3^-/-^* and *Grm6^-/-^* and their corresponding wild type littermates (respectively *Gpr179^+/+^*, *Lrit3^+/+^* and *Grm6^+/+^*) revealed 52 differently expressed genes (DEGs) in at least two of those models [14]. Metanalyses showed that half of the DEGS were associated with the term “myopia” in the literature or human genome wide association studies [14]. In addition, pathway enrichment analysis highlighted a prominent number of DEGs involved in the MAPK/ERK pathway. Among those, *Dusp4*, coding for the member of the dual specificity protein phosphatase subfamily 4 (DUSP4), drew our attention based on the literature due to its unknown function in the retina [24]. DUSP4 belongs to a group of DUSP phosphatases that inactivate their target kinases, e.g. mitogen-activated protein kinases (MAPKs), originally called extracellular signal-regulated kinases (ERKs). They are associated with cellular proliferation and differentiation by dephosphorylating both phosphoserine/threonine and phosphotyrosine residues. DUSP4 was shown to be expressed in different tissues and have a role in immunity [24], cancer [25], cardiovascular function [26], and brain [27]. *DUSP4* is expressed in the retina and more specifically in rod ON-BCs (RBCs), cone ON- and OFF-BCs, and in the retinal pigmented epithelium (RPE) [28]. It is noteworthy that induction of myopia by flickering light can be reduced by inhibiting phospho-ERK1/2 in guinea pigs [29]. In addition, retinal *Dusp4* expression appeared to be reduced upon induction of both Form Deprivation Myopia (FDM) and Lens Induced Myopia (LIM) in chicks and mice [30, 31]. Patients with a chromosomal 8p deletion, which includes the 8p12 *DUSP4*, often harbor eye defects including high myopia [32]. These findings and the expression of DUSP4 in both ON and OFF cells led us to hypothesize that the loss of DUSP4 could induce myopia through a mixed alteration of ON/OFF pathway and caused by abnormal MAPK/ERK activation. Here, we used mouse models lacking DUSP4 (*Dusp4^-/-^*) or other MAPK/ERK-associated DEGs (PKCα: *Pkcα^-/-^* and TPBG: *Tpbg^-/-^*), also downregulated in cCSNB [14] and compared them to their wild type littermates (*Dusp4^+/+^*, *Pkcα^+/+^* and *Tpbg^+/+^*). Through a combination of molecular techniques and in vivo and ex vivo recordings of ON and OFF retinal cells, we showed that the most likely light dependent function of DUSP4 in the MAPK/ERK pathway is to support emmetropization and that the absence of DUSP4 activity increases susceptibility to myopia, possibly through an altered retinal ON/OFF balance.

## Results

### Disruption of the dopaminergic pathway in *Dusp4*^-/-^, *Pkcα^-/-^*and *Tpbg^-/-^* mice

As the dopaminergic pathway is altered in several models of myopia, we sought to measure retinal levels of DA and DOPAC in *Dusp4^-/-^*, *Pkcα^-/-^* and *Tpbg^-/-^* mice. Ultra-Performance Liquid Chromatography (UPLC) measurements performed on *Dusp4^+/+^* and *Dusp4^-/-^* retinas revealed a drastic reduction of both DA (Fig. 1A) and DOPAC (Fig. 1B) in *Dusp4^-/-^* compared to *Dusp4*^+/+^ littermates (DA: 1267.3 ± 271.7 fmoles/retina in *Dusp4^-/-^*, 2467.3 ± 186.1 fmoles/retina in *Dusp4^+/+^*, Welch corrected t-test: p= 0.0270, t=3,643, df=3,539, mean difference= −1200 ± 329.4. DOPAC: 253.0 ± 10.0 fmoles/retina in *Dusp4^-/-^*, 800.3 ± 70.1 fmoles/retinas in *Dusp4^+/+^*, Welch corrected t-test: p=0.0146, t=7,725, df=2,082, mean difference= −547,3 ± 70,85). Similarly, retinal levels of DA and DOPAC were overall reduced in Pkc*α^-/-^* retinas and a trend was observed in *Tpbg^-/-^* retinas, compared to their wild-type littermates (Supplemental Figure 1A: DA: 1776,7 ± 45,9 fmoles/retina in *Pkcα^-/-^*, 2770,6 ± 109,1 fmoles/retina in *Pkcα^+/+^*, Welch corrected t-test: p=0.0053, t=8,396, df=2,689, mean difference= −994.0 ± 118.4; Supplemental Figure 1C: 2457;3 ± 50,8 fmoles/retina in *Tpbg^-/-^*, 3111,3 ± 191,9 fmoles/retina in *Tpbg^+/+^*, Welch corrected t-test: p=0.0679, t=3,294, df=2,279, mean difference= −654.0 ± 198.6. Supplemental Figure 1B: DOPAC: 444,4 ± 12,7 fmoles/retina in *Pkcα^-/-^*, 748,6 ± 89,2 fmoles/retina in *Pkcα^+/+^*, Welch corrected t-test: p=0.0733, t=3,381, df=2,081, mean difference= −304 ± 90.12. Supplemental Figure 1D: 656,0 ± 91,1 fmoles/retina in *Tpbg^-/-^*, 918,7 ± 71,9 fmoles/retina in *Tpbg^+/+^*, Welch corrected t-test: p=0.0901, t=2,262, df=3,795, mean difference= −262.7 ± 116.1). These data suggest that the lack of DUSP4, PKCα or TPBG cause an alteration of the dopaminergic pathway. It is important to note that, despite a strong difference and low variance between genotypes, quantifications of DA metabolism in the present study rely on a small sample size (n=3). Thus, statistical test was underpowered for formal hypothesis testing, and p-values should be interpreted cautiously.

**Figure 1:**
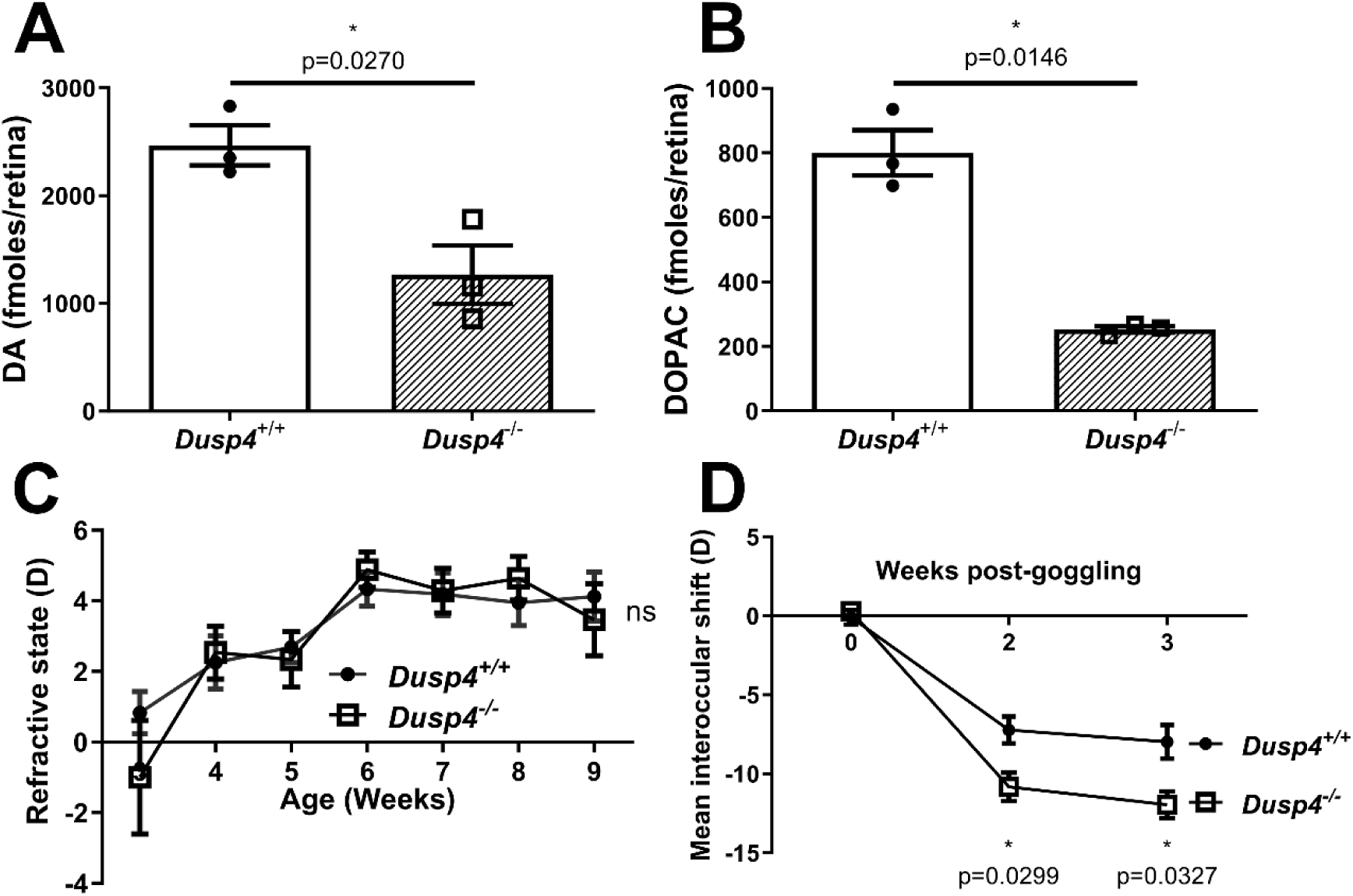
Myopic traits observed in *Dusp4^-/-^* mice. A and B: Measurement of retinal levels of DA (A) and DOPAC (B) in adult light-adapted *Dusp4^+/+^* (n=3) and *Dusp4^-/-^* (n=3) mice using UPLC. Welch t-test was used to test significance between genotype. *: p-value≤0.05. C: Measurement of spontaneous refractive development of *Dusp4^+/+^* (n=10-21) and *Dusp4^-/-^* (n=13-23) mice using infrared refractometry. Two-way ANOVA was used to test the main effect of genotype, ns: p>0.05. D: Measurement of the mean interocular shift induced by LIM in *Dusp4^+/+^* (n=9) and *Dusp4^-/-^* (n=9) mice. Two-way ANOVA was used to test the main effect of the genotype (**: p-value=0.0037). Sidak’s multiple comparison was performed as post hoc analysis for each time point, *: p≤ 0.05.

### Normal refractive development but increased sensitivity to LIM in *Dusp4^-/-^* mice

To support *Dusp4* as a candidate gene for myopia, we measured refractive development and sensitivity to LIM of *Dusp4^-/-^* mice. Measurement of the refractive state revealed that 3 weeks-old *Dusp4*^-/-^ mice were slightly-but not significantly-more myopic than *Dusp4^+/+^* littermates (Fig. 1C). From 4 weeks to 9 weeks old, *Dusp4*^-/-^ mice showed an overall normal refractive development with a hyperopic shift from 3 to 6 weeks followed by a plateau through 9 weeks of age (Fig. 1C, *Dusp4^-/-^* n=20, *Dusp4^+/+^*: n=21, RM ANOVA: main effect of genotype: ns, p=0.7248). Following LIM induction, however, *Dusp4*^-/-^ mice displayed a significantly greater myopic shift compared to *Dusp4^+/+^* littermates at 2 weeks which remained 3 weeks post goggling (Fig. 1D, −8.1±1.1 D in *Dusp4^+/+^*, n=9; −11.7±1.1 D in *Dusp4^-/-^*, n=9; main effect of genotype: RM ANOVA: p=0.0037, F (1,17) = 11,30). These results suggest that the loss of DUSP4 in mice does not affect spontaneous refractive development but increases the sensitivity to LIM.

### Expression profile and cellular immunolocalization of DUSP4 in the retina

*In situ* hybridization studies analyzed *Dusp4* mRNA at different developmental stages. At embryonic stage E15, *Dusp4* was expressed in the inner neuroblastic layer (INBL). At PN1 (PN= postnatal day), *Dusp4* was spread through both the outer neuroblastic layer (ONBL) and the INBL. From PN7 to PN66, *Dusp4* expression became more restricted to the outer part of the inner nuclear layer (INL) where BCs are located (Fig. 2A). Immunolocalization experiments confirmed BC localization of DUSP4 (Fig. 2B and 2C, green staining) in the INL, most likely in rod and cone ON-BCs. The lack of DUSP4 did not cause any change in the total DAPI+ nuclei in the INL (Supplemental figure 2A: *Dusp4^+/+^*, n= 7, in *Dusp4^-/-^*, n=9, p=0.206). Furthermore, about 30% of INL DAPI+ nuclei were stained by DUSP4 (Supplemental Figure 2B). When retinal slices from *Dusp4^+/+^* and *Dusp4^-/-^* mice were co-immun/ostained against DUSP4 (green) and against PKCα, a marker of rod BCs, or against Goα, a marker for rod and cone ON-BCs (red), BC marker immunostaining surrounded the DUSP4 positive cells (Fig. 2B and 2C). All Goα and PKCα positive cells were DUSP4 positive cells, but we observed a small subset of DUSP4 positive cells that were not stained by either anti-Goα or anti-PKCα located in the inner part of the INL. The presence of cells stained by DUSP4 but not by G0α suggest the presence of DUSP4 in a subset of OFF-BCs and/or amacrine cells (ACs). DUSP4 immunostaining was absent in retinal slices of *Dusp4*^-/-^ mice (Fig. 2B and 2C). Nonspecific immunostaining using the anti-DUSP4 antibody was observed in the outer plexiform layer in both *Dusp4^+/+^* and *Dusp4^-/-^* retinal slices. In addition, we found that the loss of DUSP4 did not induce any change in PKCα+ (Supplemental Figure 2C: *Dusp4^+/+^* retinas, n=3, *Dusp4^-/-^* retinas, n=3, p=0.400) nor in G0α+ cell number (Supplemental Figure 2D: in *Dusp4^+/+^* retinas, n=4, in *Dusp4^-/-^* retinas, n=6, p=0.3524). We found that approximately 18% of DUPS4+ cells were co-stained by PKCα and 15% of DUSP4+ cells were co-stained by G0α (Supplemental Figure 2E). Western blot analysis of retinal extracts of *Dusp4^+/+^* revealed a band at 43 kDa, consistent with the predicted molecular weight of DUSP4. This band was absent in lanes from *Dusp4^-/-^* retinal samples (Fig. 2D), validating the absence of DUSP4 in *Dusp4^-/-^* mice. Quantification of DUSP4 protein levels through western blot in P14 and P40 wild-type retinas revealed no change of DUSP4 levels during development (Fig. 2E: *Dusp4^+/+^* n= 4, *Dusp4^-/-^* n= 4, p=0.9143).

**Figure 2:**
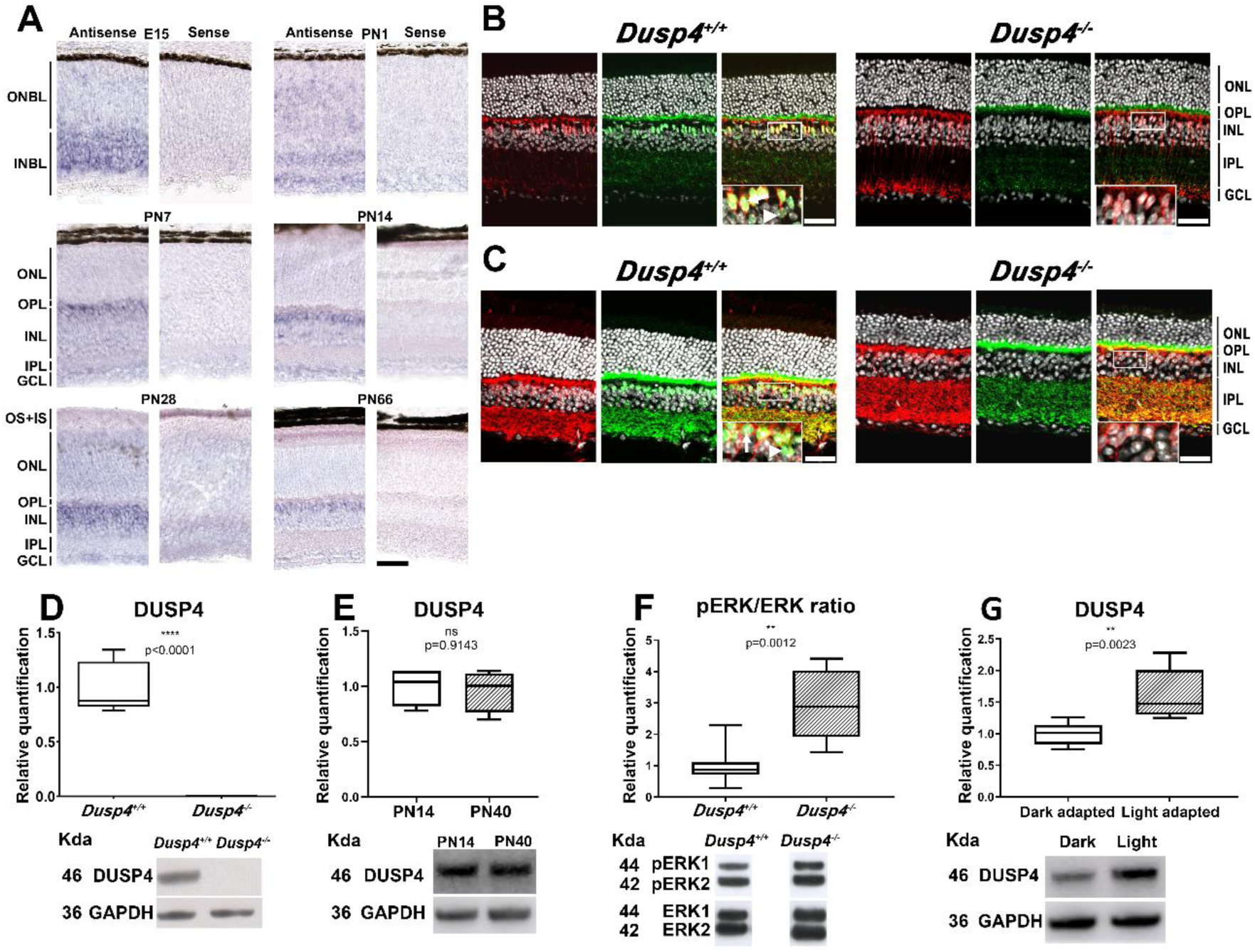
Retinal expression of DUSP4. A: ISH targeting *Dusp4* mRNA performed on wild-type retinal slices at various developmental stages (E15, PN1, PN7, PN14, PN28 and PN66). B: co-immunostaining of DUSP4 (green) with PKCα (red) performed on adult retinal slices from either *Dusp4^+/+^* (left) or *Dusp4^-/-^* (right) mice. C: co-immunostaining of DUSP4 (green) with G0α (red) performed on adult retinal slices from either *Dusp4^+/+^* (left) or *Dusp4^-/-^* (right) mice. D: Relative quantification of DUSP4 through western blot analysis performed on isolated adult retinas from either *Dusp4^+/+^* (n=5) or *Dusp4^-/-^* (n=5) mice. E: Relative quantification of DUSP4 protein through western blot analysis performed on isolated retinas from either PN14 (n=4) or PN40 (n=4) mice. F: Relative quantification of pERK/ERK ratio through western blot performed on isolated adult retinas from either *Dusp4^+/+^* (n=7) or *Dusp4^-/-^* (n=9) mice. G: Relative quantification of DUSP4 protein through western blot analysis performed on adult wild-type isolated retinas under either dark adaptation (n=6) or 50 lux light adaptation (n=7). ONBL= Outer Neuroblastic Layer, INBL= Inner Neuroblastic Layer, ONL= Outer Nuclear Layer, OPL= Outer Plexiform Layer, INL= Inner Nuclear Layer, IPL= Inner Plexiform Layer, GCL= Ganglion Cell Layer. Scale bars= 50µm. Arrowheads= Potential OFF BCs (DUSP4 positive, PKCα or G0α negative), arrows= ON-BCs (DUSP4 positive, PKCα or G0α positive). D-G: non parametric Mann-Whitney test was performed to test significance between conditions: ns: non-significant, **: P≤ 0.01, ***: P≤ 0.001. Box-and-whisker plot showing median, interquartile range, and minimum-to-maximum whiskers.

### Impact of DUSP4 loss upon the retinal ERK/pERK pathway and the impact of light

To examine whether the absence of DUSP4 alters the phosphorylation of MAPK/ERK, we performed western blot analyses with antibodies directed against ERK and its phosphorylated form: pERK (Fig. 2F). Relative to *Dusp4^+/+^* retinas (n=7), the pERK/ERK ratio was 2.98±0.37 in *Dusp4^-/-^*retinas (n=9, p=0.0012), suggesting an overactivation of the ERK pathway. These findings confirm the important role of DUSP4 in inhibiting ERK phosphorylation in the retina. We investigated whether the quantity of DUSP4 is influenced by light. Although DUSP4 is detectable under dark conditions, DUSP4 levels were increased at 50 lux light-adapted retinas (n=7) compared to dark-adapted retinas (n=6) (Fig. 2G: p=0.0023), suggesting that light can increase DUSP4 expression.

### Cellular localization of DUSP4 loss in cCSNB mouse models

As previously shown, *Dusp4* transcripts were reduced in cCSNB mouse models, *Gpr179^-/-^* and *Lrit3^-/-^* [14]. Here, we confirmed the reduction of *DUSP4* mRNA (Fig. 3A, n=6 for each group, p=0.0084 for *Gpr179^-/-^*, p=0.0031 for *Lrit3^-/-^*) and protein levels (Fig. 3B, n=5 for each group, p=0.0317 for *Gpr179^-/-^*, p=0.0079 for *Lrit3^-/-^*) in cCSNB retinas. Using immunostaining, we observed that the number of cells labelled by DUSP4 was reduced by half in both *Gpr179^-/-^* (Supplemental Fig. 3A, n=11 p<0.0001) and *Lrit3^-/-^* (Supplemental Fig. 3A, n=10 p<0.0001) INLs. To determine the cellular population directly affected by the reduction ofDUSP4 in cCSNB mouse models, we performed co-immunostaining against DUSP4 with either Secretagogin (SCGNB), an EF-Hand Calcium Binding Protein and a marker of cone ON and OFF BCs, or ISLET1, a marker of all ON-BCs, ACs and retinal ganglion cells (RGCs) in adult *Lrit3^+/+^*, *Lrit3^-/-^, Gpr179^+/+^* and *Gpr179^-/--^* retinal slices (Fig. 3C to 3F and Supplemental Fig. 3A to 3G). No change was observed in the number of SCGN+ nor ISLET1+ cells in *Gpr179^-/-^* and *Lrit3^-/-^* INL (Supplemental figures 3B and 3C, n=5-6 for each group). Quantification of cells co-stained by DUSP4 and SCGN unveiled an overall unchanged proportion of DUSP4+SCGN+ cells nor DUSP4+SCGN-in any of the cCSNB mouse models tested (Supplemental Figure 3D and 3E, n=5-6 for each group). On the other hand, quantification of cells co-stained by DUSP4 and ISLET1 highlighted a drastic decrease in the proportion of cells DUSP4+ISLET1+ in both *Gpr179^-/-^* (n=5-6 for each group, p=0.0043) and *Lrit3^-/-^*(n=5 for each group, p=0.0317) INLs and a corresponding increase in the proportion of cells DUSP4+ISLET1-(Supplemental Fig. 3F and 3G). The micrographs shown in Fig. 3C to 3F reveal that the loss of DUSP4 staining in cCSNB retinas mainly imply the most outer cells of the INL. These results suggest the decrease of DUSP4 expression in *Gpr179^-/-^* and *Lrit3^-/-^* retinas in most present in both rod-BCs and cone ON-BCs, while DUSP4 expression in cCSNB OFF-BCs remains unchanged.

**Figure 3:**
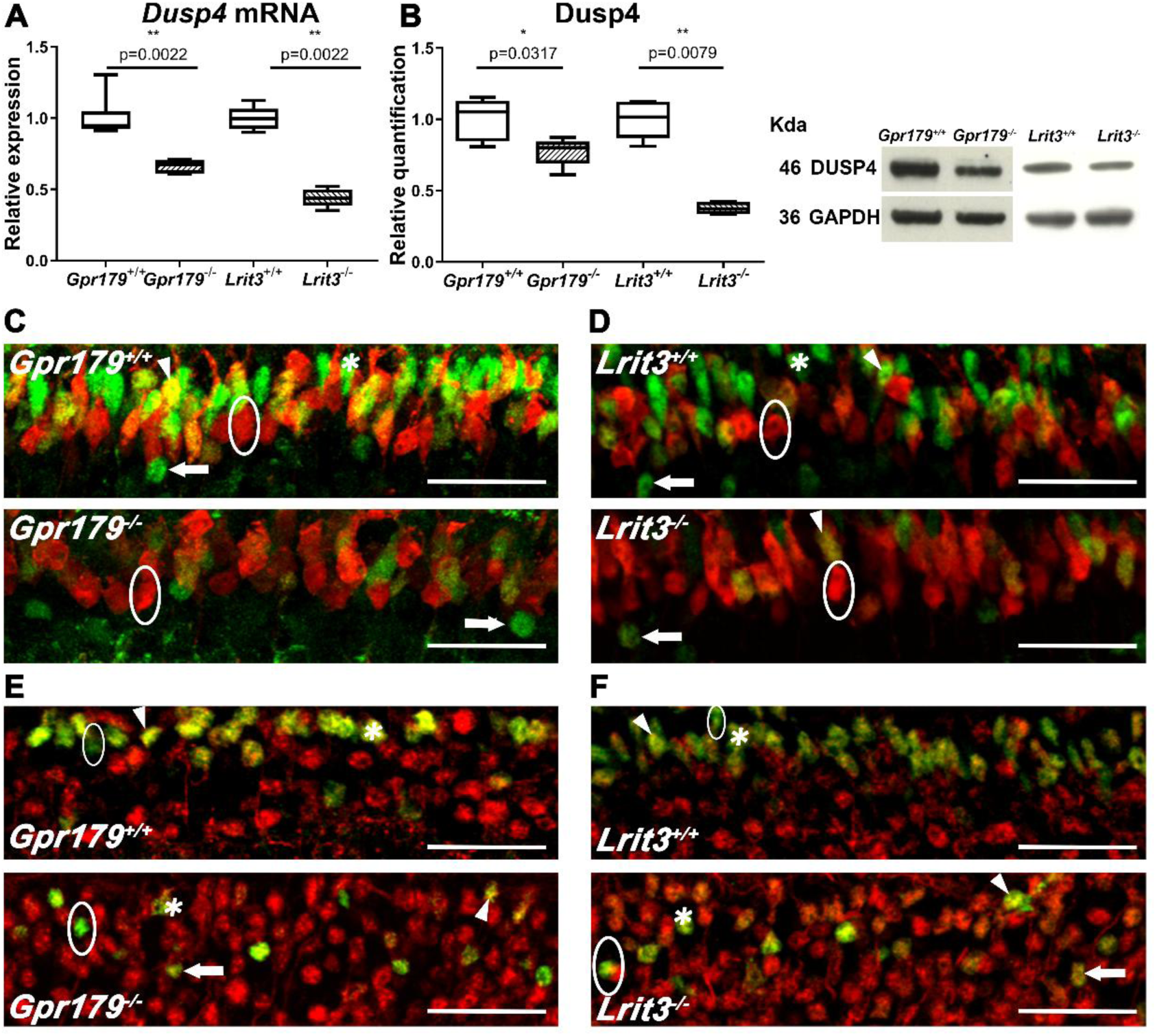
Impact of ON-pathway activity upon DUSP4 expression. A: Quantification of *Dusp4* mRNA by RT-qPCR performed on isolated adult retinas from two cCSNB mouse models (*Gpr179^-/-^* and *Lrit3^-/^*, n=6 each*^-^*) and their corresponding wild-type littermates (n=6 each). B: Quantification of DUSP4 protein by western blotting performed on isolated adult retinas from two cCSNB mouse models (*Gpr179^-/-^* and *Lrit3^-/-^*, n=5 each) and their corresponding wild-type littermates (n=5 each). C and D: Co-immunostaining of DUSP4 (green) with SCGN (red) performed on adult retinal slices from either *Gpr179^-/-^* (C, lower image) or *Lrit3^-/-^* (D, lower image) and their corresponding wild-type littermates (upper images). E and F: Co-immunostaining of DUSP4 (green) with ISLET1 (red) performed on adult retinal slices from either *Gpr179^-/-^* (E, lower image) or *Lrit3^-/-^*(F, lower image) and their corresponding littermates (upper images). Scale bars= 50µm. Arrowheads= cone ON-BCs (DUSP4 positive, SCGN or ISLET1 positive, upper part of the INL), stars= RBCs (DUSP4 positive or negative, SCGN negative, ISLET1 positive, upper part of the INL), arrows= potential ACs (DUSP4 positive, SCGN negative, ISLET1 positive, lower part of the INL), circles= potential OFF-BCs (SCGN positive, ISLET1 negative, DUSP4 positive or negative, lower part of the INL). Non parametric Mann-Whitney test was used to test the significance between genotypes: *: P≤ 0.05, **: p-value≤ 0.01.

### Impact of DUSP4 loss on mouse visual acuity

To investigate whether the loss of DUSP4 leads to altered visual acuity, optomotor tests were performed with increasing spatial frequencies upon 6 week- and 6-month-old *Dusp4*^+/+^ and *Dusp4*^-/-^ mice under both photopic and scotopic conditions. Under both scotopic and photopic conditions, optomotor responses were unchanged in 6 weeks-old (Figs. 4A, n=9-11, p=0.9413 and B, n=8 for each group, RM ANOVA: p=0.4505), but reduced in 6-month-old *Dusp4^-/-^* mice (Figs. 4C, n=10-12, RM ANOVA: p=0.0041 F (1, 20) = 10,49, Sidak’s multiple comparison: p=0.0093, t= 2,560, DF= 17,33 for scotopic cpd= 0.063 and p<0.0001, t= 3,406, DF= 18,15 for scotopic cpd=0.125) and 4D, n=10-12, p=0.1449, Sidak’s multiple comparison: p=0.0145, t= 3,051, DF= 100,0, for photopic cpd=0.063) compared to *Dusp4*^+/+^ age-matched littermates at the lowest spatial frequencies. The effect at 6 months was greater for scotopic than photopic responses. We did not observe any change in the visual acuity. These data suggest that the deletion of *Dusp4* causes an impairment of the optomotor response, more marked under scotopic conditions, and possibly in an age-dependent manner.

**Figure 4:**
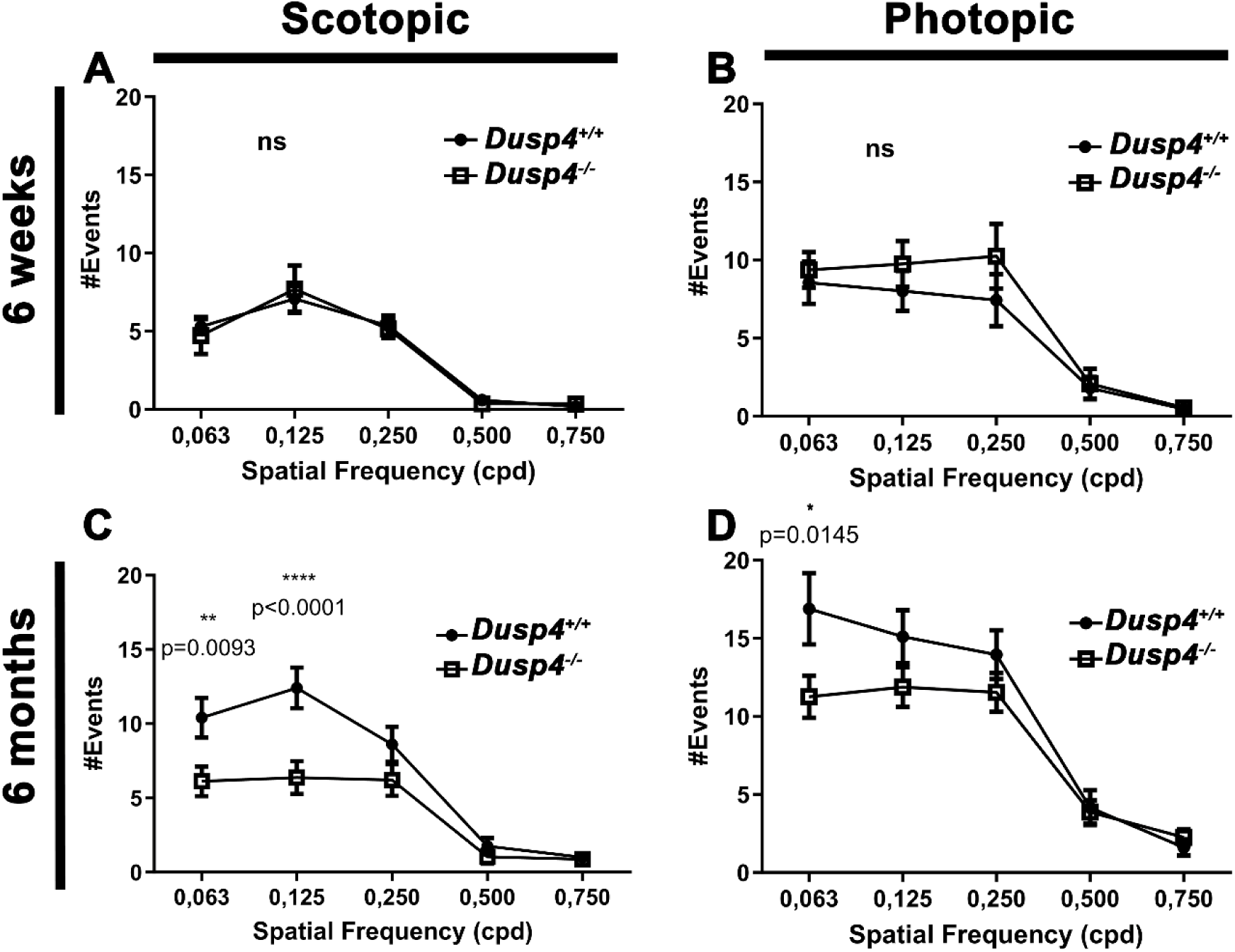
Altered optomotor response in *Dusp4^-/-^* mice. A and B: Number of positive events obtained with 6-week-old *Dusp4^-/-^* mice and their wild-type littermates under scotopic (A, n=9-11) and photopic (B, n=8 each) conditions. C and D: Number of positive events obtained with 6-month-old *Dusp4^-/-^* mice and their wild-type littermates under scotopic (C, n=10-12) and photopic (D, n=10-12) conditions. Cpd= cycles per degree. Repeated measures ANOVA was used to test the main effect of the genotype: A-B: p≥0.05, C: p≤0.001, D: p≤0.05. Sidak’s multiple comparison was performed as post hoc analysis to compare differences for each spatial frequency: *: P≤ 0.05, **:P≤ 0.01 ***: P≤ 0.001.

### Retinal architecture in mice lacking DUSP4

As functional disturbances can be correlated with morphological alterations, optical coherence tomography and eye fundus examinations were used to determine whether structural alterations could be observed in *Dusp4^-/-^* retinas (see Supplemental Figure 2 for an example with measurements of INL and IPL thickness, n=5-6 for each group). No obvious morphological alterations were observed in *Dusp4^-/-^* retinas at either 6 weeks or 6 months of age, suggesting that *Dusp4* ablation does not cause gross morphological changes.

### Alteration of retinal signaling in mice lacking DUSP4

In order to determine whether the genetic ablation of *Dusp4* could cause detectable alterations of retinal signaling, we performed full-field electroretinogram (ERG) recordings on 6 week- and 6-month-old *Dusp4^+/+^* and *Dusp4^-/-^* mice (Fig. 5) and multiple electrode array (MEA) analyses on isolated retinas from 6-month-old *Dusp4^+/+^* and *Dusp4^-/-^* mice (Fig. 6). In 6-week-old *Dusp4^+/+^* and *Dusp4^-/-^* mice, no difference was observed in any of the ERG parameters tested (Fig. 5C: RM ANOVA: n=8-9, p=0.3393, 5D: RM ANOVA: n=8-9, p=0.6790, 5E: RM ANOVA: n=8-9, p=0.7948, 5F: RM ANOVA: n=8-9, p=0.4946). Fig. 5A and 5B show representative traces obtained at −1.5 log cd.s/m² (rod responses) and at 1.5 log cd.s/m² (mixed rod-cone responses). At 6 months of age, the amplitude of the b-wave was significantly increased in *Dusp4^-/-^* mice compared to *Dusp4^+/+^* (Fig. 5K: RM ANOVA: n=9-10, p=0.0186, F (1, 17) = 6,771, Sidak’s multiple comparison: p=0.0414, t= 3.170, DF= 15,81 for 0.5log cd.s/m²; p=0.0210, t= 3.477, DF= 16.37 for log cd.s/m².). No statistical change in the a-wave amplitude (Fig. 5I: n=9-10, RM ANOVA: p=0.1727) was observed. The implicit time of both a wave (Fig. 5J: n=9-10, RM ANOVA: p=0.0976) and b-wave (Fig. 5L: n=9-10, RM ANOVA: p=0.6168) was unchanged. Representative traces are shown in Fig. 5G and 5H.

**Figure 5:**
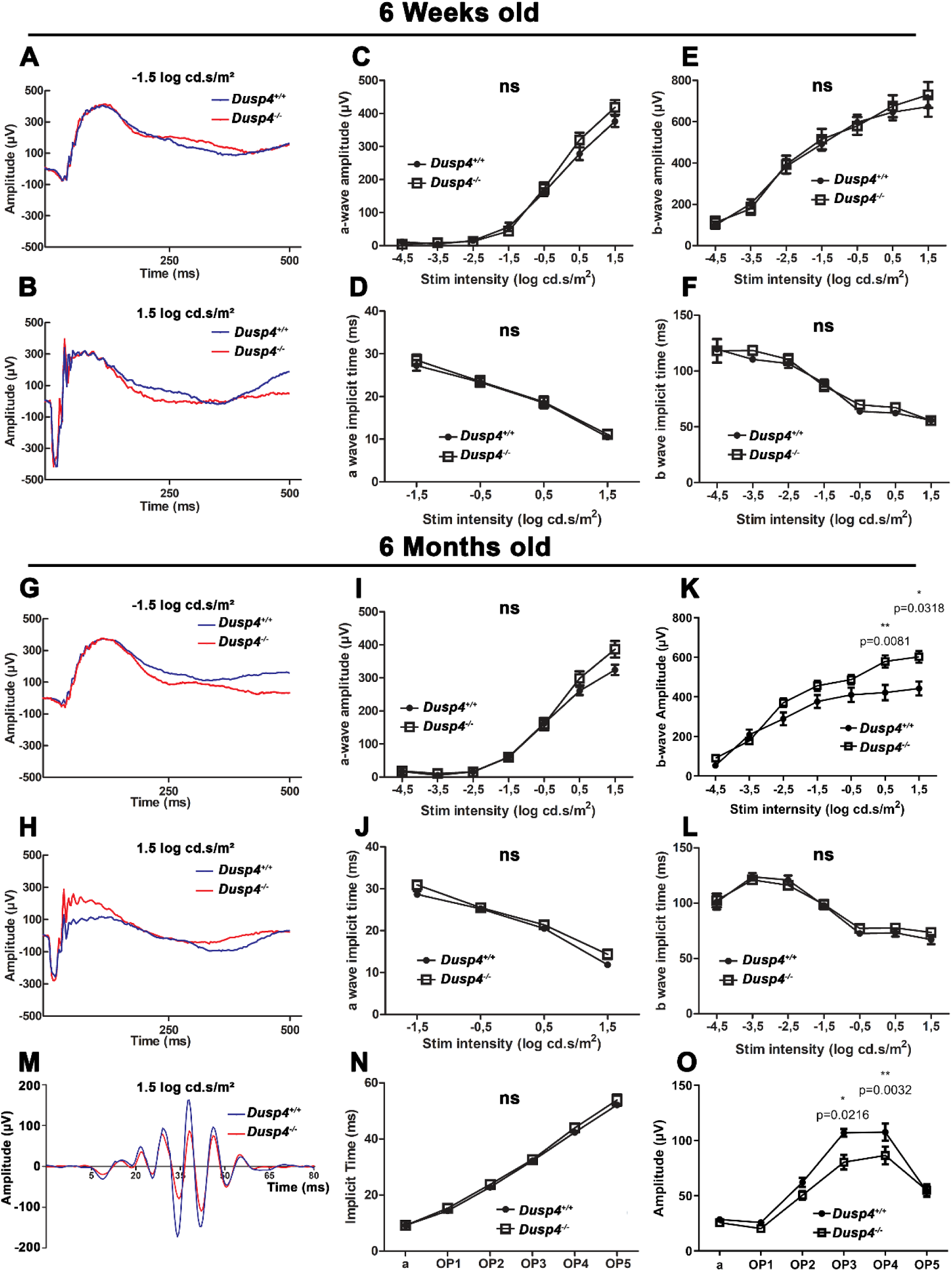
Altered b-wave and OPs observed in *Dusp4^-/-^* mice. A to F: Measurement of the amplitude (C and E) and implicit time (D and F) of the ERG a-wave and b-wave in response to flashes of increasing intensities recorded from 6-week-old *Dusp4^-/-^* (n=9) and their wild-type littermates (n=8). A and B show representatives ERG traces at −1.5 log cd.s/m² and 1.5 log cd.s/m², respectively. G to L: Measurement of the amplitude (I and K) and implicit time (J and L) of a-wave and b-wave in response to flashes of increasing intensities recorded from 6-month-old *Dusp4^-/-^* (n=10) and their wild-type littermates (n=9). G and H show representatives ERG traces at −1.5 log cd.s/m² and 1.5 log cd.s/m², respectively. N to O: Measurement of implicit time (N) and amplitudes (O) of the a-wave (x axis: a) and the OPs (x axis: OP1 to OP5) elicited by flashes of 1.5 log cd.s/m² from 6-month-old *Dusp4^-/-^* mice (n=11) and their wild-type littermates (n=13). A to O: Repeated measure ANOVA was used to test main effect of genotype. Sidak’s multiple comparison was performed as post hoc analysis to compare differences for each stimulus intensity, ns: p>0.05 *: p≤0.05.

**Figure 6:**
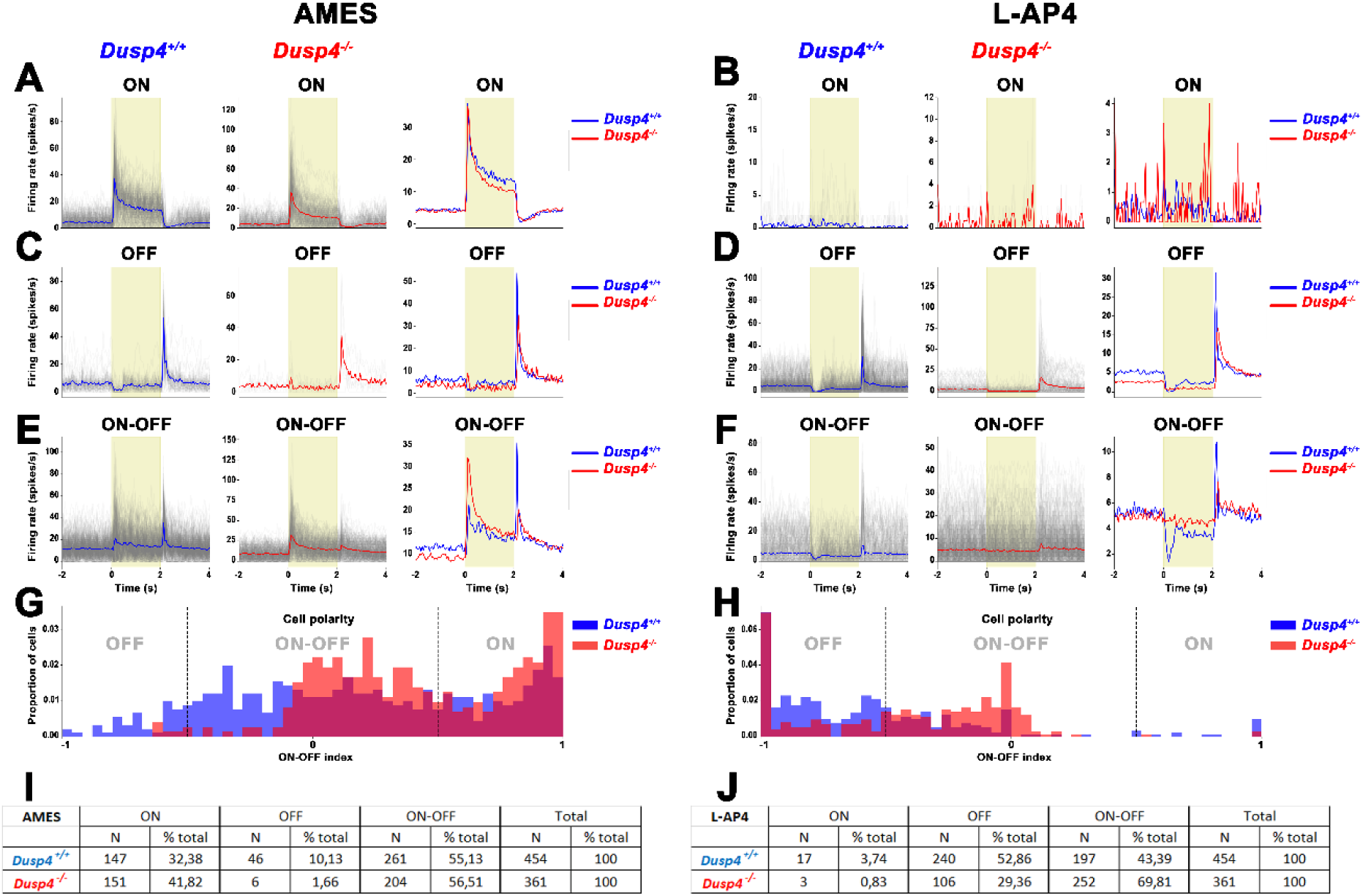
Altered ON and OFF RGCs signaling in *Dusp4^-/-^* mice. Mean spiking rate of ON-RGCs (A: 147 *Dusp4^+/+^* cells, 151 *Dusp4^-/-^* cells and B: 17 *Dusp4^+/+^* cells, 3 *Dusp4^-/-^* cells), OFF RGCs (C: 46 *Dusp4^+/+^* cells, 6 *Dusp4^-/-^* cells and D: 240 *Dusp4^+/+^* cells, 106 *Dusp4^-/-^* cells), ON-OFF RGCs (E: 261 *Dusp4^+/+^* cells, 204 *Dusp4^-/-^* cells and F: 197 *Dusp4^+/+^* cells, 252 *Dusp4^-/-^* cells), distribution of RGCs (G and H) according to their response to light onset (ON) or offset (OFF) or both (ON-OFF) and summary of the number and proportion of recorded cells for each condition (I and J) in isolated *Dusp4^+/+^* (blue, n=2) and *Dusp4^-/-^* (red, n=2) retinas under normal AMES (left) or L-AP4 (right) perfusion. Yellow parts represent the light stimulus. Non parametric Mann-Whitney test was used to test significance of the genotype upon ON-OFF index in recorded cell population.

Furthermore, the amplitudes of the scotopic oscillatory potentials (OPs) of *Dusp4^-/-^* mice were significantly decreased at 6 months (Fig. 5O: n=11-13, RM ANOVA: main effect of genotype: p=0.0021, F (1, 25) = 11,80, Sidak’s multiple comparison: p=0.0135, t= 3,548, DF= 18,24 for OP3, p= 0.0166, t= 3,317, DF= 25,00 for OP4) without any change in their implicit time (Fig. 5N: n=11-13, p>0.9999. Representative traces are shown in Fig. 5M). The change in OPs suggests that *Dusp4* loss alters retinal signaling mediated by ACs, causing an improper response to light stimuli under scotopic conditions.

Full-field MEA recordings performed on dark-adapted 6-month-old *Dusp4^-/-^* retinas perfused with a normal recording medium (AMES) unveiled an unchanged firing rate of ON-RGCs in response to light onset (Fig. 6A) but a markedly reduced firing rate of OFF-RGCs in response to light offset (Fig. 6C). Notably, the ON-OFF index was significantly higher (p=2.02*10^-12^) in *Dusp4^-/-^* (n= 361 cells in 2 animals, ON-OFF index= 0.427704± 0.02056) retinas compared to *Dusp4^+/+^* (n= 454 cells in 2 animals, ON-OFF index= 0.17788± 0.02408) as a low number of *Dusp4^-/-^* OFF-RGCs were responsive (Fig. 6C, 6G and 6I). An increased ON-response, but reduced OFF-response of *Dusp4^-/-^* ON/OFF-RGCs was observed in response to light onset and offset, respectively (Fig. 6E), compared to *Dusp4*^+/+^ mice. In addition, both ON- and OFF-responses of *Dusp4^-/-^* ON/OFF-RGCs showed a slightly delayed return to baseline activity after onset of the corresponding stimulus (Fig. 6E). Under pharmacological blockade of the ON-pathway through L-AP4 perfusion, almost no ON-activity was recorded from both genotypes (Fig. 6B), as expected. A reduced spiking rate of *Dusp4^-/-^* OFF-RGCs was still observed, but a higher number of *Dusp4^-/-^* OFF-RGCs were responsive to the stimulus (Fig. 6D, 6H and 6J). Similarly, the ON-OFF index was higher (p=4.42*10^-12^) in *Dusp4^-/-^* retinas (ON-OFF index= −0.410828± 0.02091, n= 361 cells in 2 animals) compared to *Dusp4^+/+^* retinas (ON-OFF index=-0.580639 ± 0.01976, n=464 cells in 2 animals). The spiking rate of the OFF-response of *Dusp4^-/-^* ON/OFF-RGCs was increased in the presence of L-AP4 (Fig. 6F) and a higher number of ON/OFF-RGCs were responsive, compared to wild-type (Fig. 6H and 6J). These data suggest that the loss of *Dusp4* induces an imbalance between ON- and OFF-pathway activity at the level of RGCs, which is most noticeable in OFF- and ON/OFF-RGCs.

## Discussion

Despite a growing global awareness and improved medical care, the prevalence of myopia is increasing worldwide. Thus, further studies are required to gain a better understanding of myopia development and progression. Previous work [16–19] including our [12–14] converged toward a predominant implication of ON pathway upon myopia onset. Here, we show that reduced retinal DA and sensitivity to myopia can occur despite higher ON-BC and lower OFF-RGC responses, potentially caused by altered MAPK/ERK activation in several retinal cell types. We highlight here a complex interplay between retinal ON, OFF pathways and emmetropization.

### Effects of *Dusp4* upon retinal DA and ocular response to myopic defocus

The reduced levels of retinal DA and the higher sensitivity of *Dusp4^-/-^* mice to LIM validated *Dusp4* as a myopia-associated gene. Despite these findings, *Dusp4^-/-^* mice also show a higher scotopic b-wave amplitude, reduced OPs amplitude and reduced OFF responses at the ganglion cell level as opposed to the complete loss of b-wave and OPs observed in cCSNB models with relatively preserved OFF-responses on ganglion cell levels [33, 34]. These findings suggest that an alternative chain of events is leading to myopia in *Dusp4^-/-^* eyes compared to cCSNB models and exclude the candidacy of *Dusp4* as a cCSNB gene. In addition, our findings indicate that Dusp4 does not substantially alter baseline refractive development, but it increases vulnerability to lens-induced myopia. This phenotype supports a gene–environment interaction in which DUSP4 contributes to the retinal mechanisms that determine how visual experience is translated into ocular growth signals. Thus, *Dusp4* may be considered as a susceptibility gene that becomes critical under myopia-inducing conditions such as defocus rather than as a normal refractive development factor.

### DUSP4 localization, time course and light modalities of expression

DUSP4 expression in other biological tissues is regulated by several factors including growth serum, oxidative stress, and UV light [26, 35, 36]. Our experiments add to these findings as retinal DUSP4 expression appears to be affected by light. In addition, mice lacking ON-pathway activity show reduced DUSP4 expression in ON-BCs, with unchanged DUSP4 immunostaining in ACs or OFF-BCs (Fig. 3). Taken together, these findings suggest that 1) circadian rhythm, with changes in ambient light, may impact DUSP4 expression and 2) increased *Dusp4* expression in light-adapted retinas requires a functional ON-pathway. A recent study reported reduced *Dusp4* expression in retinas from chick occluded eyes [37] in keeping with statement 1). Furthermore, it is conceivable that the remaining DUSP4 staining in dark adapted eyes is limited to OFF-BCs.

Pronounced *Dusp4* mRNA staining in the INBL during embryonic stages and in INL at post-natal stages is consistent with previous analyses showing *Dusp4* expression in BCs in the adult retina [14, 38] as the INBL gives rise to RGCs, ACs and BCs [39]. According to databases [38] and our Immunolocalization study with specific markers, DUSP4 is not restricted to ON-BCs (RBCs and BC6 ON-CBC subtype) but is also expressed in some OFF-BCs (potentially BC2 and BC1B subtypes). With the present findings, we cannot exclude that DUSP4 is localized in ACs, but to our knowledge, this is not supported by transcriptomic and protein databases [40]. This complexifies the interpretation of the role of *Dusp4* in the retina and pinpoints the need for investigations dissecting the specific role of DUSP4 in these cell types.

### Potential role of DUSP4 on Direction sensitive ganglion cells (DSGCs)

Our phenotyping experiments unveiled that *Dusp4^-/-^* mice have an overall lower optomotor response without any change in visual acuity. Our experimental paradigm implies a rotating velocity of about 12°/sec. At this velocity, ON-OFF DSGCs are known to be the major actors responsible for optomotor behavior [41–43]. Implication of *Dusp4* in ON-OFF DSGCs is supported by its expression in BC2 and BC1B CBCs, as they contact ON-OFF DSGCs, and the altered ON-OFF RGCs response observed in our MEA experiments. Altered ON-OFF DSGCs are not known to have any mechanistical relationship with myopia onset, thus, it is possible that our optomotor data highlight an role for DUSP4 in retinal direction-selective circuit, independent from myopia.

### Altered scotopic ERG in mice lacking DUSP4

Consistent with previous findings [44], the amplitude of the ERG is age-dependent, irrespective of the presence of DUSP4. Nevertheless, its deletion leads to increased scotopic b-wave amplitude associated with reduced OPs at the highest stimulus intensities, strongly suggesting that the absence of DUSP4 causes an increased depolarization of ON-BCs in response to light. However, at these light intensities, both ON and OFF pathways contribute to the b-wave amplitude, in a positive and negative manner, respectively. Thus, as *Dusp4* is expressed in some subtypes of OFF-BCs, we cannot exclude the implication of the OFF pathway in the increased b-wave through a lack of hyperpolarization possibly causing a delayed return to the baseline. Due to the excitatory nature of the ON-BC to DAC synapse, releasing glutamate during light conditions, it is expected that a higher b-wave amplitude will increase rather than decrease DA release as observed in *Dusp4^-/-^* retinas. This discrepancy can be explained by the fact that neuronal depolarization does not necessarily imply higher neurotransmitter, such as glutamate release and that pERK/ERK ratio have a complex impact on synaptic activity.

### Putative MAPK/ERK-mediated mechanisms of synaptic transmission and sensitivity to myopic defocus

The role of MAPK/ERK signaling in BCs function remains incompletely defined, but evidences from other neuronal systems provide a useful framework for interpretation. In cortical and hippocampal neurons, synaptic pERK/ERK levels bidirectionally regulate neurotransmitter release [45–49]. Transient, physiological increases in pERK/ERK can enhance release through a synapsin 1–dependent mechanism [50], whereas sustained elevations, typical of pathological conditions, are associated with reduced synaptic output [51]. Prolonged ERK activation may recruit phosphatases such as calcineurin, limiting phosphorylation of synapsin 1 and thereby impairing vesicle mobilization. In parallel, ERK signaling may also constrain Ca²⁺ entry: inhibition of ERK1/2 phosphorylation increases neurotransmitter release via recruitment of L-type calcium channels [52, 53], suggesting that pERK can negatively regulate Ca²⁺ influx, potentially through channel sequestration.

In the context of our data, these observations support the hypothesis that elevated and sustained pERK/ERK levels in *Dusp4^-/-^* retinas reduce glutamate release at the ON-BC to amacrine cell synapse, leading to decreased DA release. This interpretation is consistent with phenotypes observed in other mouse models linked to MAPK/ERK regulation. Notably, Pkcα^-/-^ and *Tpbg^-/-^* mice, both associated with dysregulation of this pathway and downregulated in cCSNB models [14], display similar ERG alterations, reduced retinal DA levels (Supplemental Figure 1), and are suspected to have an increased susceptibility to myopia [54–56]. TPBG, in particular, has been implicated in vesicle exocytosis, and its loss reduces glutamate release from ON-BCs to ACs, causing reduced OPs despite an increased scotopic b-wave amplitude [47]. Together, these findings raise the possibility that DUSP4, PRKCA, and TPBG converge on shared signaling mechanisms involving MAPK/ERK-dependent phosphorylation of presynaptic substrates such as synapsin 1, thereby modulating synaptic transmission.

Altogether, these data and our findings suggest that a precise balance of MAPK/ERK activation and deactivation in BCs may be critical for maintaining normal retinal output and downstream dopaminergic signaling, with potential implications for refractive development. However, several limitations remain. First, the increased pERK/ERK ratio was measured in whole retina, preventing cell type–specific attribution to ON-versus OFF-BCs. Second, ERK-dependent effects on neurotransmitter release are thought to require presynaptic localization of phosphorylated ERK, but the subcellular distribution of pERK in Dusp4⁻/⁻ retinas is unknown. Addressing these points as well as synapsin 1 and L-type calcium channel will be essential to refine the proposed mechanism and its link to myopia susceptibility.

The altered OFF and ON/OFF activity observed in our *ex vivo* recordings from *Dusp4^-/-^*RGCs is consistent with the known expression of *Dusp4* in both ON- and OFF-BCs [38]. It is noteworthy that the stimulation parameters used here allow for the stimulation of intrinsically photosensitive RGCs (ipRGCs). The flash duration corresponds to the bare minimum of mouse ipRGCs time to response (2 seconds) [57]. As our trains of stimulation consist in 2 seconds flashes followed by 10 seconds of darkness, we cannot rule out the possibility that ipRGCs were stimulated here. Nevertheless, L-AP4 perfusion completely abolished ON-RGCs activity, suggesting that the impact of ipRGCs activation was neglectable in our case. In addition, the criteria we used to compute the ON, OFF, ON-OFF index of RGCs prevents our analysis from considering ipRGCs population. Thus, we cannot exclude that DUSP4 deficiency can impact ipRGCs activity, but our MEA paradigm most likely prevented them from being considered in our analysis. Deciphering the specific impact of DUSP4 upon ipRGCs remains of interest for future studies.

The potential mechanism through which the loss of DUSP4 causes sensitivity to myopia involves the dysregulation of MAPK/ERK activation (for a summary of the proposed mechanisms, see figure 7). We hypothesize that the absence of DUSP4 in ON-BCs induces a strong and sustained pERK/ERK ratio, which reduces glutamate release at the RBC to A2 amacrine cells synapse and at the ON-CBC to DAC synapse through a synapsin 1 and/or L-type calcium channel mechanism. This causes a lower activity of ACs, highlighted by the lower OPs amplitude and the reduced retinal levels of dopamine metabolism. The decreased AC activity leads to lower inhibitory feedback to ON-BCs, observed through the increased scotopic b-wave amplitude. The unchanged ON-RGC activity observed during our ex vivo recordings can reflect the paired alterations of ON-BCs and ACs which both influence ON-RGCs activity. It is widely accepted that activation of the ON-pathway is independent of OFF-pathway activity, while activation of the OFF-pathway implies both an intrinsic activation of OFF-cells and a reduction of inhibition from the ON-pathway, mediated by inhibitory ACs [58, 59]. As L-AP4 infusion allows an incomplete recovery of the number of recordable OFF-RGCs, we can suggest that the altered *Dusp4^-/-^* OFF-RGC response imply both a higher inhibition from the ON pathway and a lower activation by OFF-CBCs. Finally, the altered ON and OFF responses observed in *Dusp4^-/-^* ON/OFF-RGCs is also evidence of the dysregulation of both ON and OFF pathway in *Dusp4^-/-^* mice.

**Figure 7:**
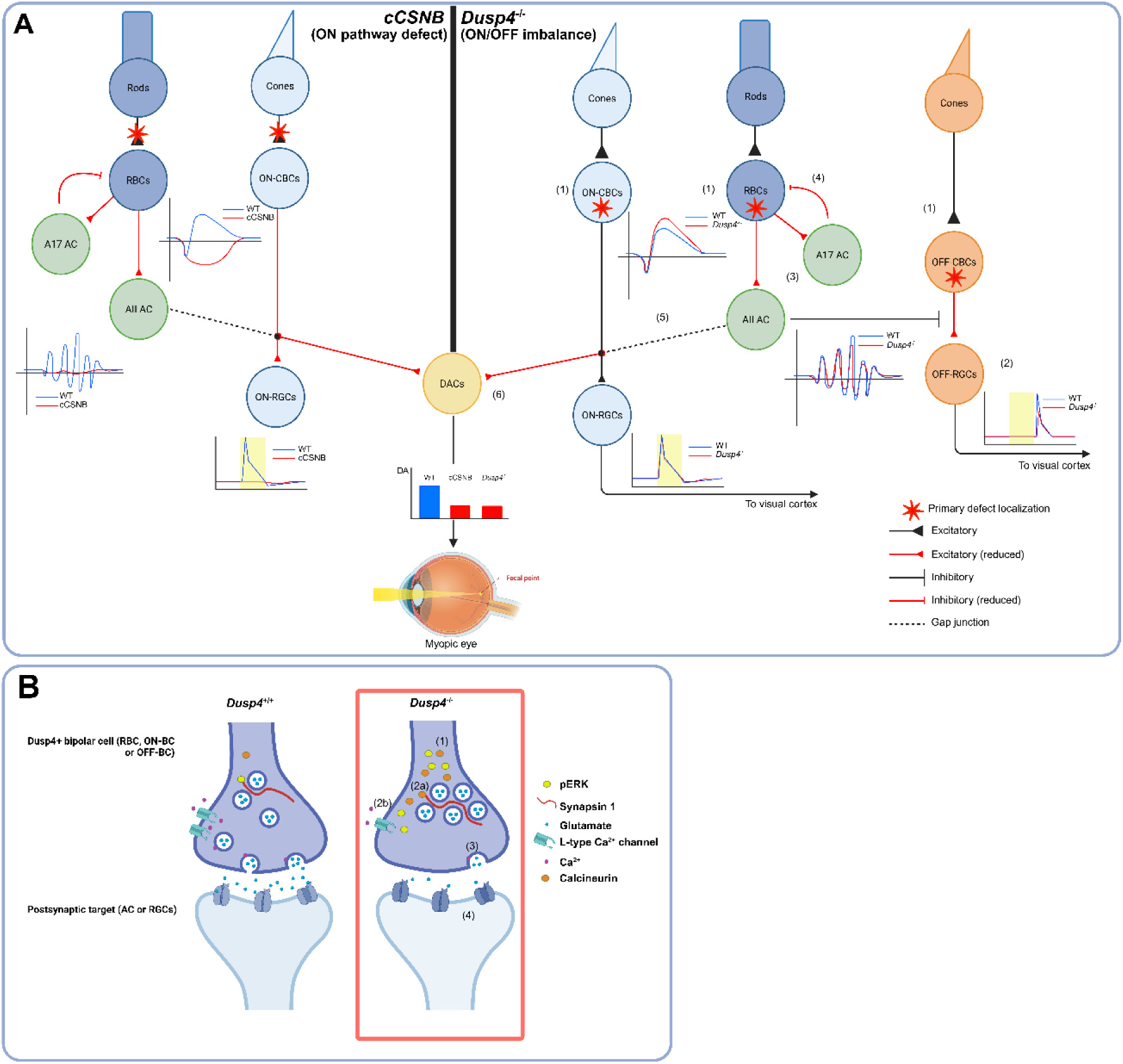
Proposed mechanisms leading to myopia and altered retinal signaling caused by DUSP4 deficiency compared to cCSNB. Left: genetic disruption of one of the cCSNB-causing gene leads tom the complete loss of transmission between photoreceptors and ON-BCs (red stars), observed through an electronegative ERG. This negatively impacts ACs activity and remove the excitatory output upon DACs, reducing DA release and ultimately causing abnormal ocular elongation. Right: alternative chain of events occurring in *Dusp4^-/-^* retinas. A: imbalance of ON-OFF activities in *Dusp4^-/-^* retinas. pERK/ERK is increased in all retinal cells depleted from DUSP4 (RBCs, ON-CBCs and OFF-CBCs), causing a reduction of neurotransmitter release from those cells (1). The decrease of OFF-RGCs activity, observed during *ex vivo* MEA recordings (2), arises from a combination of lower excitatory input from OFF-CBCs and higher inhibition from the ON-pathway. The lower, but not abolished, amplitude of OPs, observed during scotopic ERGs, is meaningful of a reduced activity or synchronicity of amacrine cells, most probably those connected to BCs (DACs, A17, A2). Amacrine AII and A17 ACs receive less excitatory input from RBCs (3), the higher b-wave observed during ERGs comes from reduced reciprocal feedback from A17 Acs (4). The decreased glutamate release at the RBC-A2 synapse ultimately causes a reduction of DACs activity through the A2-ACs to ON-CBCs gap junction (5) (dashed line). Ultimately, DACs release less DA, reducing the STOP signal cascade that normally prevents abnormal eye growth (6). B: Proposed mechanisms for the reduction of neurotransmitter release caused by the loss of DUSP4. The absence of DUSP4 leads to high and sustained levels of pERK in all bipolar cells (1). Under normal conditions, pERK phosphorylates synapsin 1 at SER4/5 site. When abnormally abundant, pERK recruits large amount of phosphatases such as calcineurin which prevent the phosphorylation of SER4/5 sites (2a), reducing the amount of readily releasable pool of synaptic vesicles (3). In parallel, large amount of pERK can reduce the quantity of L-type calcium channels present at the synapse by promoting their sequestration (2b), causing a reduction of Ca^2+^ influx normally arising from neuronal depolarization. Intrasynaptic deficiency of Ca^2+^ leads to reduced fusion of vesicles with membrane at the synaptic cleft. Both 2a and 2b mechanisms can occur independently and do not exclude each other. Ultimately, this results in decreased glutamate release, leading to lowered activation of postsynaptic receptors, and, consequently, reduced stimulation of postsynaptic neurons such as amacrine and ganglion cells. Created in BioRender. Navarro, J. (2026) (URL: https://BioRender.com/o1sj4lu)

An important limitation must be noted as DUSP4 is constitutively deprived in our model. LIM and retinal DA are known to be mediated by local ocular mechanisms solely, but as *Dusp4* is expressed in RPE, and as RPE is known to participate in ocular response to defocus and eye growth [60, 61], we cannot exclude a partial participation of RPE-related mechanisms in the development of myopic traits. Dissecting the specific role of *Dusp4* in the RPE and its putative impact on axial elongation is of interest for future studies. In addition, *Dusp4* is expressed in the brain (https://www.proteinatlas.org/ENSG00000120875-DUSP4/tissue). It was previously shown that *Dusp4^-/-^* mice have reduced synaptic plasticity in the hippocampus [53]. Hippocampal plasticity is thought to be a major component of spatial memory which may also contribute to the phenotype observed upon optomotor response recordings in *Dusp4^-/-^* mice as on any behavioral assessment. Therefore, optomotor response should be confirmed in future studies by using a retina specific knock out of *Dusp4*.

Altogether, the present study and our previous work [14] confirm *Dusp4* as a gene implicated in the development of myopia. The present study highlights MAPK/ERK pathway as a potential therapeutic target to control syndromic and non-syndromic myopia. In accordance with previous findings obtained from *Aplp2^-/-^* mice and *GJD2* mutations in myopic patients, we report that sensitivity to myopia does not imply solely ON-pathway defect but can also occur through an imbalance of ON and OFF pathways. Future studies should focus on challenging the hypothesis of reduced glutamate release by bipolar cells through hyperactivation of MAPK/ERK signaling. Similarly, assessing other myopia-associated DEGs found in cCSNB [14] and implicated in MAPK/ERK pathway, such as *Prkca*, *Tpbg* or *Ptprr* would be relevant. Validating their impact on pERK/ERK, RGCs activity and myopia onset in mouse models and in patients could help to bridge the gap between altered MAPK activity, ON/OFF imbalance and myopia onset. In addition, further dissection of the retinal and RPE role of *Dusp4* by using cell type-specific deletion methods could be insightful to obtain a deeper understanding of retinal signaling and its implication in ocular elongation.

## Materials and Methods

### Animal Care and ethical statement

We have complied with all relevant ethical regulations for animal use. Animal procedures were performed according to the Council Directive 2010/63EU of the European Parliament, the Council of September 22, 2010, and on the ARRIVE 2.0 guidelines on the protection of animals used for scientific purposes. They were approved by the French Minister of National Education, Superior Education and Research. Mouse lines and projects were registered as following: *Dusp4^-/-^*: APAFIS#22074 *Gpr179*^-/-^: APAFIS#18037, and *Lrit3*^-/-^: APAFIS#18132, refractometry: APAFIS #27474.). We used both male and female mice. They were kept in 12:12 hour light:dark cycles with mouse chow and water ad libitum. The environmental enrichment consisted of cotton nests and cellulose sticks. The MGI codes for each mouse lines are as follow: *Dusp4^-/-^*: MGI:2442191, *Gpr179^-/-^*: MGI:2443409, *Lrit3^-/-^*: MGI:2685267, *Tpbg^-/-^*: MGI:5007391, *Pkcα^-/-^*: MGI: 97595. The origins of mouse lines, including supplier information are referenced as follow: *Dusp4^-/-^*: [24]:, *Gpr179^-/-^*: [34], *Lrit3^-/-^*[33]:, *Tpbg*^-/-^: [62], *Pkcα*-/-: [63]. The corresponding genotyping protocols are referenced as follow: *Dusp4^-/-^*: see below, *Gpr179^-/-^* [34], *Lrit3^-/-^* [33]:, *Tpbg^-/-^* [62]:, *Pkcα^-/-^*[63]. The *Dusp4^-/-^* and *Gpr179^-/-^* mouse lines originate from pure C57Bl6/N genetic background. *Tpbg^-/-^*, *Pkcα^-/-^* and *Lrit3^-/-^* mouse lines all originate from mixed 129/SvEv-C57BL/6 genetic background. The corresponding controls all consist in wild-type littermates.

### Polymerase chain reaction (PCR) genotyping for *Dusp4*

DNA was extracted from mouse tails with 50 mM NaOH after incubation at 95°C for 30 min. Wild-type and mutant allele were amplified independently using a polymerase (HOT FIREPol, Solis Biodyne, Tartu, Estonia), two specific forward primers: m*Dusp4*_3F (5’-GTCTAGAAAGCCTTTCCGTC-3’) for the wild-type allele and m*Dusp4*_F3 (5’-GATCCGATAACTTCGTATAGC-3’) for the mutant allele and two specific reverse primers: m*Dusp4*_3R (5’-GCGCCTAACGCCTAAAATCC-3’) for the wild-type allele and m*Dusp4*_4R (5’-GTAAAACAGGAGCTGGCATC-3’) for the mutant allele. The following PCR program was used: 15 min at 95°C for denaturation, 35 cycles of: 45 sec at 95°C, 1 min at 58°C, and 1 min 30 sec at 72°C, and for final extension, 10 min at 72°C. This gives rise to the following amplicons: PCR using m*Dusp4*_3F and m*Dusp4*_3R primers amplifies a product of 386 base pairs (bp) for the wild-type allele and no product for the mutant allele, PCR using m*Dusp4*_F3 and m*Dusp4*_4R primers amplifies no product for the wild-type allele and a 1000 bp product for the mutant allele. PCR products were separated by electrophoresis on 2% agarose gels, stained with ethidium bromide, and visualized using a documentation system (Molecular Imager^®^ Gel Doc^TM^ XR+ System, Bio-Rad, Hercules, California, USA).

### Induction and measurement of myopia in *Dusp4^-/-^* mice Lens induced myopia

The lens induced myopia protocol was adapted from a previously validated protocol [64]. Briefly, P21 mice were anesthetized by isoflurane inhalation (5% induction, 2% maintenance) during the whole procedure. The scalp was cut to expose 1.2 cm of the skull in the rostrocaudal axis. Two intracranial screws were implanted on both left and right sides of the skull at y = −2 mm from the bregma. A homemade goggle frame was placed on the skull and fixed using dental cement (FujiCEM™, Phymep© Cat #900903, Paris, France). The goggle frame, adapted from [45], was built in resin using a 3D printer. −25 D lenses were stuck on the frame using glue (vetbond™, Phymep©, Paris, France). Stitches were used to avoid displacement of lens by mice. The −25 D lens was always placed in front of the right eye for three weeks. Every two days, the lenses were removed for cleaning and the eyes were rehydrated using Lacrifluid 0.13% eyedrops.

### Refractometry

After a dark adaptation period of 30 min, eye drops were used to dilate the pupils (0.5% mydriaticum, 5% neosynephrine). The mice were placed on a homemade restraining platform in front of an eccentric infrared photorefractometer; its calibration was described elsewhere [65]. The mouse was positioned so that the first Purkinje image was in the center of the pupil. The data were then recorded using software designed by Franck Schaeffel [65]. We collected two data sets per animal, one per eye, each data set consisting of 90 measures, each of which being the mean of 10 successful measures. For refractive development experiments, the measurements were performed once per week, every week from 3 to 9 weeks old and the mean between both eyes was used for statistical analysis. To evaluate the sensitivity to myopia induction, the difference between goggled and ungoggled eyes measured at P21 (day of surgical procedure), P35 (14 days after surgery) and P42 (21 days after surgery) was used for statistical analysis. Only the non-myopic mice displaying less than 2D of interocular shift at P21 were used for the induction protocol.

### DA and DOPAC measurements

P80 Mice were euthanized by CO2 inhalation followed by cervical dislocation at noon. The light adaptation and the measurement of retinal levels of DA and DOPAC were performed as previously described [12].

### Western blot quantification of DUSP4 protein and pERK/ERK ratio Preparation of retinas for protein extraction for western blot analyses

P42 Mice were euthanized by CO2 inhalation followed by cervical dislocation at noon. Whole retina samples were lysed with lysis buffer (Tris 50 mM pH7.5, NaCl 150 mM, Triton X100 1%) for 30 min on ice, vortexed every 10 min. Samples were thereafter sonicated 3×10 sec, with 30 sec on ice between each sonication. Soluble proteins were recovered from the supernatant following a 10-min centrifugation (13,000 rpm) at 4°C. The concentration of protein in each sample was determined by a bicinchoninic acid (BCA) assay and samples were kept at −80°C until required.

### Western blot analyses

Equal concentrations (6μg) of protein extracted from whole retina of wild-type and *Dusp4^-/-^* mice, were separated by electrophoresis on a commercially available 1 mm protein gel (NuPAGE^TM^ 4-12% Bis-Tris Gel, Invitrogen, Waltham, MA, USA). The proteins were transferred to a 0.2 μm nitrocellulose membrane (Trans-Blot Turbo Mini Nitrocellulose membrane, Bio-Rad, Hercules, CA, USA) using a semi-dry transfer system (Trans-Blot^®^ Turbo, Bio-Rad) and blocked in 5% dry milk (Regilait, Saint-Martin-Belle-Roche, France) in phosphate-buffered saline containing 0.05% Tween-20 (PBST) at room temperature (RT) for 1 hour. Membranes were incubated with the primary antibodies consecutively at 4°C overnight. Description of the primary antibodies, their dilution and references can be found in Supplemental table 1. Subsequently, the membranes were washed three times with PBST and incubated with the corresponding anti-rabbit, −mouse or -goat horseradish peroxidase (HRP) secondary antibodies (1/10,000) for 1 hour at RT. After three washes in PBST, membranes were incubated with enhanced chemiluminescence (ECL) substrate for 5 min. Antibody binding was thereafter detected in a dark room using a commercially available developer (Kodak, Rochester, NY, USA).

### Protein quantification

Protein quantification was done using western blot images and by a commercial software (ImageJ, National Institutes of Health, Bethesda, MD, USA).

### Preparation of retinal sections for localization of *Dusp4* protein and RNA

#### Preparation of retinal sections

Retinal slices of 6-week-old *Dusp4^-/-^*, *Gpr179^-/-^* and *Lrit3^-/-^* mice and their corresponding wild-type controls were obtained according to a protocol adapted from [66]. Briefly, P42 Mice were euthanized by CO2 inhalation followed by cervical dislocation. For hybridization in situ (ISH) entire eyes were isolated, a hole was performed just behind the ora serrata and then eye was fixed in 4% paraformaldehyde (PFA) at RT during 20 minutes. After fixation the eye was washed three times in PBS and put directly on 30% cold sucrose solution for the cryoprotection. For immunolocalization studies the anterior segment and lens were removed, and the eyecup was fixed in cold 4% PFA for 20 minutes. After three washing with cold PBS, eyecups were cryoprotected with increasing concentrations of sucrose (10% and 20% for 1h each and 30% overnight).

To finish, the entire eye for ISH or the eyecups for immunolocalization were embedded in 7.5% gelatin 10% sucrose and frozen in a dry ice-cooled isopentane bath. Sections were cut at a thickness of 20 µm on a cryostat (model HM560, Microm; Thermo Fisher, Walldorf, Germany). and mounted onto glass slides (Super-Frost plus Gold for ISH and Super-Frost plus for immunolocalization, ThermoFisher Scientific,Waltham, MA, USA). The slides were air dried and stored at 80 °C

#### In situ hybridization procedure

RNA in situ hybridization was performed on adult mouse retinal section (20µM). The mouse cDNA corresponding to *DUSP4* exon 1-4 was cloned in a plasmid pBluescript II SK by Genecust company (Genecust, Boynes, France). The antisense and sense probes were obtained after plasmid was linearized using the restriction enzymes NotI and KpnI. Antisense and sense RNA in situ hybridization probes were synthesized using T7 and T3 RNA polymerase (Roche Diagnostics, Basel, Switzerland), respectively, and labeled with digoxigenin-UTP (Roche Diagnostics). Mouse retinal sections were postfixed 10 minutes in 4% PFA, washed with PBS, and treated with proteinase K (10mg/mL; Invitrogen, Carlsbad, Czech Republic) for 2 minutes. Following a wash with PBS, the sections were postfixed in 4% PFA, washed in PBS, and then acetylated in an acetylation buffer of 1.3% triethanolamine (Sigma-Aldrich, St. Quentin Fallavier, France), 0.25% acetic anhydride (Sigma-Aldrich), and 0.06% hydrochloric acid solution. The sections were washed in 1% Triton X-100 PBS solution and blocked for 2 hours in hybridization buffer containing 50% formamide, 5X SSC (saline sodium citrate [1X SSC is 0.15 M NaCl plus 0.015 M sodium citrate]), 1X blocking solution (Denhardt’s solution; Sigma-Aldrich), 250 µg/mL yeast tRNA (Roche Diagnostics), and 240 µg/mL salmon sperm (Roche Diagnostics), pH 7.4. The sections were hybridized with digoxigenin-labeled probes overnight at 72°C, after which they were rinsed for 2 hours in 0.2% SSC at 72°C and blocked for 1 hour at RT in 0.1 M Tris, 0.15 M NaCl (B1), 10% normal goat serum (NGS; Vectorshield, Burlingame, CA), pH 7.5. After blocking, slides were incubated overnight at RT with anti-digoxigenin antibody conjugated with alkaline phosphatase (1:5000 dilution; Roche Diagnostics) in B1 containing 1% NGS. After additional washes, the alkaline phosphatase activity was detected using nitro blue tetrazolium chloride (337.5 mg/ml; Roche Diagnostics) and 5-bromo-4-chloro-3-indolyl phosphate (175 lg/ml; Roche Diagnostics). Five hours later, sections were mounted in Mowiol (Calbiochem, Merck, Carlstadt, NJ). Slides were scanned with a Nanozoomer 2.0 high throughput (HT) equipped with a 3-charge–coupled device time delay integration (TDI) camera (Hamamatsu Photonics, Hamamatsu, Japan).

#### Immunohistochemistry procedure

P42 Mice were euthanized by CO2 inhalation followed by cervical dislocation at noon. Retinal slices of *Dusp4^-/-^*, *Gpr179^-/-^* and *Lrit3^-/-^* mice and their corresponding wild-type littermates were placed in a humid chamber. Slices were saturated using a PBSGT solution (PBS1X, 0.2% gelatin, 0.25% TritonX100) overnight at RT. Incubation with primary antibodies diluted in PBSGT lasted overnight at RT. Description of primary antibodies can be found in supplemental table 1. After washing in PBS 0.1% TritonX100, the sections were incubated with corresponding secondary antibodies coupled to Alexa Fluor 488 or Alexa Fluor 594 (Jackson Immuno Research Laboratories, Baltimore, MD, USA) at a dilution of 1:1000 in PBSGT for 1h at RT. The slides were stained with 40,6-diamidino-2-phenylindole (DAPI) sections were washed 3 times in PBS and mounted with cover-slipped with mowiol (Calbiochem, Merck, Carlstadt, NJ). Images were acquired using a confocal microscopy (Olympus FV1200, Rungis, France). Cellular populations stained by markers (Dapi, DUSP4, G0α, PKCα, SCGN or ISLET1) were manually quantified using the cell counter plugin from ImageJ (National Institutes of Health, Bethesda, MD, USA).

#### Quantification of Dusp4 mRNA Preparation of retinas for RNA extraction

P42 Mice were euthanized by CO2 inhalation followed by cervical dislocation at noon. RNA extraction of whole retina samples was performed according to manufacturer’s instructions (RNeasy Protect Mini Kit, Qiagen, Hilden, Germany). The Dnase elimination step was included and followed according to the manufacturers’ protocol. A final volume of 40 µl was eluted using Rnase-free water. The RNA yield was analyzed via spectrophotometric absorbance (Nanodrop^TM^ 2000, Thermo Fisher Scientific, Waltham, MA, USA), with suitable concentration being above 200 ng/µl. Extracted RNA was kept at −80°C until required.

#### Real Time PCR (RT-PCR)

RT-PCR was performed on extracted RNA from whole mouse retinas. The quality of RNA in each sample was first tested on a bioanalyzer (Agilent 2100 Bioanalyzer, Agilent Technologies, Santa Clara, CA, USA) according to manufacturer’s instructions (Agilent RNA 6000 Nano Kit, Agilent). Samples with a RIN <7 were discarded. A two-step RT-PCR was then performed using RNAs with concentrations of 50 ng/μl for each sample and 5 ng/μl for the negative controls. RNAs were reverse transcribed into single-stranded cDNA according to the kit protocol (QuantiTect^®^ Reverse Transcription, Qiagen). The cDNA was first diluted to a concentration of 2 ng/μl for each sample and 0,5 ng/μl for the negative control samples. The PCR reaction solutions each contained 5μl cDNA, 10 μl SYBR Green, 0,4μl of each primer (Table 2) and 4,2 μl H2O, for a total reaction volume of 20 μl. Each RT-PCR reaction was performed in triplicates for both the wild type (wt) and the mutant mouse, with glyceraldehyde-3-phosphate dehydrogenase (GAPDH) used as a reference gene. The detailed protocol of the PCR-conditions will be delivered on request.

**Table 1:** description of antibodies used in the present study.

| Antibody | Size (kDa) | Dilution | Host | Reference | Manufacturer |
| --- | --- | --- | --- | --- | --- |
| DUSP4 | 43 | 1/500 | Rabbit | ab216576 | Abcam |
| GAPDH | 36 | 1/200 | Goat | SC-20357 | Santa Cruz |
| PKCα | 80 | 1/500 | Mouse | P5704 | Sigma-Aldrich |
| G0α | 40 | 1/200 | Mouse | MAB3073 | Merck Millipore |
| ISLET1 | 39 | 1/500 | Mouse | Ab86501 | Abcam |
| SCGN | 32 | 1/100 | Mouse | AMAb90630 | Sigma Aldrich |
| PERK | 42,44 | 1/1000 | Rabbit | 9101S | Cell Signaling |
| ERK | 42,44 | 1/1000 | Rabbit | 4695T | Cell Signaling |

**Table 2:**
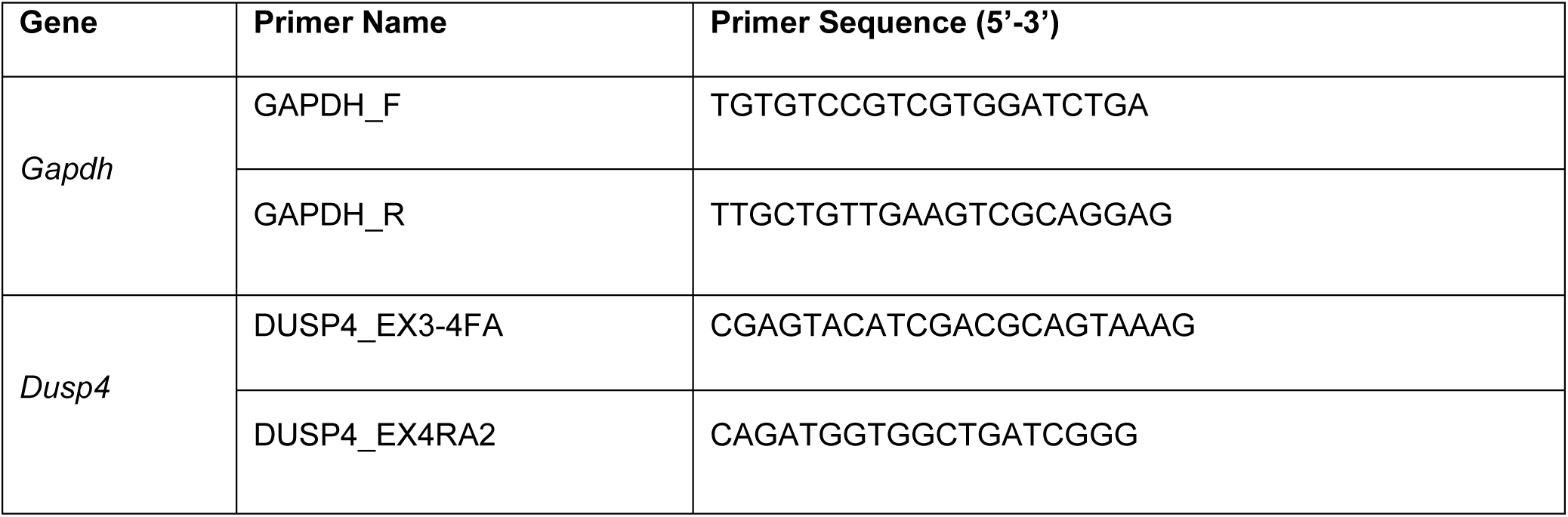
primers used in the present study.

#### Phenotyping of *Dusp4^-/-^* mice Optomotor response

6-week-old or 6-month-old mice were dark-adapted overnight before the optomotor test. Mice were placed on a grid platform at the center of a motorized drum (adapted from [67]) revolving clockwise and anticlockwise at two revolutions per minute and covered by vertical black and white stripes of a defined spatial frequency (0.063, 0.125, 0.25, 0.5 and 0.75 cycles per degree (cpd)). Each test was recorded with a digital infrared camera to count head movements of the mice. Head movements in both directions were considered to obtain the number of events per minute. The experimenter responsible for analysis of the videos was blinded for the genotype of mice.

#### Electroretinography

After overnight dark adaptation, 6-week-old or 6-month-old mice were anesthetized with intraperitoneal injection of a mixture consisting of ketamine (80mg/kg, Axience, France) and xylazine (8mg/kg, Bayer HealthCare, France) diluted in sterile 0.9% NaCl. Eye drops were used to dilate the pupils (0.5% mydriaticum, Thea, France and 5% neosynephrine, CSP, France) and anesthetize the cornea (0.4% oxybuprocaïne chlorhydrate, Thea, France). Upper and lower lids were retracted to keep eyes open and bulging. Contact lens electrodes for mice (Mayo Corporation, Japan) were placed on the corneal surface to record ERG responses through a layer of Lubrithal (Dechra Veterinary Products, Denmark) and impedance was controlled as it was not higher than 10MΩ. Needle electrodes placed subcutaneously in forehead and in the back served as reference and ground electrodes. Stimulus presentation and data acquisition were provided by the Espion E2 system (Diagnosys LLC, Lowell, MA, USA). For the dark adapted ERG recordings, a cut-off filter of 0.312-300Hz (low-high) was applied and eight levels of stimulus intensity ranging from 0.0006 cd.s/m^2^ to 60 cd.s/m^2^ were used. Each scotopic ERG response represents the average of five responses from a set of five flashes of 4ms stimulation. Oscillatory potentials were recorded under dark condition. A cut-off filter of 75-300Hz (low-high) was applied to the signal and an average of five responses to a set of five flashes of 4ms stimulation at 60 cd.s/m^2^ was obtained. The mean between the two eyes was considered as the experimental unit for analysis. Mice showing an interocular difference of the response amplitude at the highest scotopic stimulus intensity (>20% of the maximum) were excluded from analysis.

#### Spectral Domain-Optical Coherence Tomography (SD-OCT)

6-week- or 6-month-old mice were anesthetized by isoflurane inhalation (5% for induction, 2% to maintain) during the whole procedure. Eye drops were used to dilate the pupils (0.5% mydriaticum, 5% neosynephrine). SD-OCT images were recorded for both eyes using a spectral domain ophthalmic imaging system (Bioptigen, Inc., Durham, NC, USA). We performed rectangular scans consisting of a 1.4 mm by 1.4 mm perimeter with 1000 A-scans per B-scan with a total B-scan amount of 100. Scans were obtained first while centered on the optic nerve, and then with the nerve displaced either temporally/nasally or superiorly/inferiorly. We measured the thickness of Retinal Nerve Fiber Layer and Ganglion cell layer together (RNFL+GCL), Inner plexiform Layer (IPL), Inner Nuclear Layer (INL), Outer Plexiform Layer (OPL), Outer Nuclear Layer (ONL), External Limiting Membrane (EML), Inner Segment and Outer Segment together (IS+OS) and Retinal Pigment Epithelium and Choroid together (RPE+C).

#### Assessment of *Dusp4^-/-^* retinal circuitry using multi-electrode array (MEA)

Tissue preparation, and stimulation protocol were performed as previously described [68]. After overnight dark adaptation, 6 months old mice were euthanized by CO_2_ inhalation followed by cervical dislocation. Retinas were carefully dissected under dim-red light and conserved in Ames medium (Sigma-Aldrich, St. Louis, MO, USA) oxygenated with 95% oxygen and 5% CO_2_. Retinas were placed on a Spectra/Por membrane (Spectrum Laboratories, Rancho Dominguez, CA, USA) previously coated with poly-D-lysine and pressed against an MEA (MEA256 100/30 iR-ITO; Multi Channel Systems MCS, Reutlingen, Germany) using a micromanipulator, RGCs facing the electrodes. Retinas were continuously perfused with bubbled Ames medium or L-AP4 (50µM, Tocris, Bristol, UK) at 34.8°C at a rate of 2 ml/min and let to rest for 45 min before the recording session. Under dark conditions, 10 repeated full-field light stimuli at a 450 nm wavelength were applied to the samples at 4.85^13 photons/cm^2^/s for 2 seconds with 10-seconds interval by using a Polychrome V monochromator (Olympus, Hamburg, Germany) driven by an STG2008 stimulus generator (MCS). Raw RGC activity recorded by MEA was amplified (gain 1000–1200) and sampled at 20 kHz using MCRack software (MCS).

The resulting data was high-pass filtered at 100 Hz, and spikes were detected using SpyKING CIRCUS [69]. Only cells presenting a low number of refractory period violations (inferior to 0.5%, 2ms refractory period) and well-distinguished template waveforms were selected for analysis, giving a total of 887 cells (526 from two *Dusp4^+/+^* mice and 361 from two *Dusp4^-/-^* mice). These criteria ensured the high quality of the reconstructed spike trains. Subsequent analysis was performed using custom Python scripts.

We determined for each sorted RGC the maximum firing frequency in an interval of 2 seconds after light onset (for ON-responses) and in an interval of 2 seconds after light offset (for OFF-responses). These values were normalized to the mean spontaneous firing frequency of the corresponding RGC. Considering that significant responses have a maximum firing frequency that is superior to the mean spontaneous firing frequency +5 SD, we determined the time at which these significant frequencies were reached after the light onset for ON-responses and after the light offset for OFF-responses. We computed an ON-OFF index I_ON-OFF_ for each cell with the following formula: *I_ON−OFF_ = (n_light_ − n_dark_) ÷ (n_light_ + n_dark_)* with n_light_ the number of detected spikes during the light onset and n_dark_ those detected for the same duration after the light offset. Cells having a I_ON-OFF_ close to −1 would be considered OFF cells, while those having a I_ON-OFF_ close to 1 would be ON cells. Here we considered cells as OFF with I_ON-OFF_<-0.5, ON with I_ON-OFF_>0.5, and ON-OFF in between.

#### Statistics and reproducibility

The experimental unit consists in individual animals. In dopamine measurements, spontaneous refractive development, WB and RT-qPCR quantifications, ERG recordings, OCT quantifications, and optomotor response assessment, means of the two eyes for each animal were considered as one experimental unit. For sensitivity to myopia induction, the difference between the two eyes is considered as the experimental unit. For IHC quantifications and MEA *ex vivo* recordings, only one eye per animal, chosen randomly, was used as experimental unit. The sampling size was determined according to previously peer-reviewed studies with similar experimental designs. All the statistical tests used here are Two-tailed. We applied an *a priori* exclusion criteria for the lens-induced myopia experiments: mice with spontaneous myopic refractive state and/or an interocular shift >2D before surgery were excluded from the experiment. Values are all presented as mean ± SEM. Significance was tested by non-parametric tests when normality could not be assumed. The statistical tests, potential post hoc analysis, p-values significance and other statistics components can be found in figures legends and in results section. Main effect of genotype is mentioned in main text. Post-hoc analysis are mentioned in figures and captions. All statistical tests were performed using Graphpad (Prism 10.0.2, for Windows, GraphPad Software, Boston, Massachusetts USA, www.graphpad.com).

## Data availability

All raw data used for analysis are provided in a repository (https://doi.org/10.5281/zenodo.19881381). The full code used for MEA is referenced [69].

## Supporting information

Supplemental figures, legends and uncropped WB gels

## Fundings and awknowledgmens

Supported by Agence Nationale de la Recherche (ANR-22-CHIN-0006) (OM, CZ); Banque publique d’investissement (PREMYOM) (OM, CZ), Retina France (CZ); Valentin Haüy and AFM (CZ), IRPINSERM (CZ and RD); Prix Dalloz pour la recherche en ophtalmologie and Fondation Dalloz—Institut de France (CZ); Fondation Voir et Entendre (CZ); Fondation de l’oeil—Fondation de France (IA, CZ), Ville de Paris and Region Ile de France; Labex Lifesenses (reference ANR-10-LABX-65) (IA, CZ), supported by French state funds managed by the ANR within the Investissements d’Avenir programme (ANR-11-IDEX-0004-0); the Programme Investissements d’Avenir IHU FOReSIGHT (ANR-18-IAHU-01) (IA, CZ); National Institutes of Health grant EY029985 (RD), funded by the European Union under Grant Agreement #101119501 (SB). The authors are grateful to Manuel Simonutti, Julie Dégardin, Pauline Abgrall and Marion Cornebois for their valuable help in p²henotyping (platform at the Institut de la Vision). The authors are thankful to Julie Geus, Raja Rabia, Wissam Afyass and Lisa-Léa Rota for the mouse handling and housing (Janvier labs®).

