## Supplemental figures, legends and uncropped WB gels for "Increased sensitivity to myopia and altered retinal ON/OFF balance in a mouse model lacking *Dusp4*"

### Supplemental figures and table.

This file contains:

List of Myotreat participants

Supplemental figures 1 to 4

Supplemental table 1

Legends for Supplemental figures 1 to 4

Legend for Supplemental table 1

Uncropped blots for Figures 2D, 2E, 2F, 2G and 3B

Legends for uncropped blots

**The Myotreat consortium**:

PIs:

Baraas Rigmor,

Feldkaemper Marita,

Guggenheim Jeremy,

Klaver Caroline C.W.,

Schroedl Falk,

Wahl Siegfried,

Zeitz Christina

DCs:

Alihodzic Arjana,

Boranijasevic Sanja,

Castells Nieto Anna Sofia,

Cid Vinas Pol,

He Xi,

Jannat Juli Farjina,

Konwar Jahnobi,

Mishra Mayank,

Neto Rodrigues Ana Miguel,

Nguyen Thu-Nga,

Spanic Filip,

Stauffer Kee Lucien

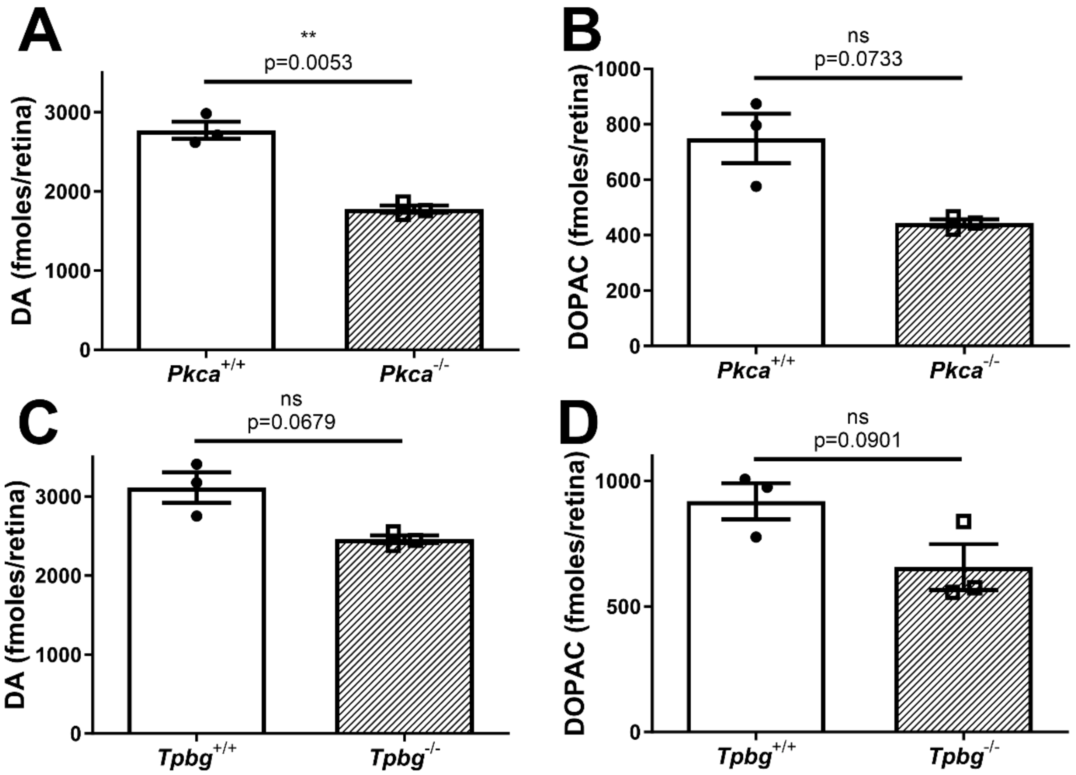


**Supplemental figure 1: Reduced levels of DA and DOPAC in *Pkcα*$^{-/-}$ and *Tpbg*$^{-/-}$ retinas.**

Measurement of retinal levels of DA (left panels) and DOPAC (right panels) in adult light-adapted *Pkcα*$^{-/-}$, *Pkcα*$^{+/+}$ (upper panels) and *Tpbg*$^{-/-}$, *Tpbg*$^{+/+}$ (lower panels) mice using UPLC (n=3 each). Welch t-test was used to test significance between genotypes. ns= non-significant, p-value>0.05; **: p-value ≤0.01.

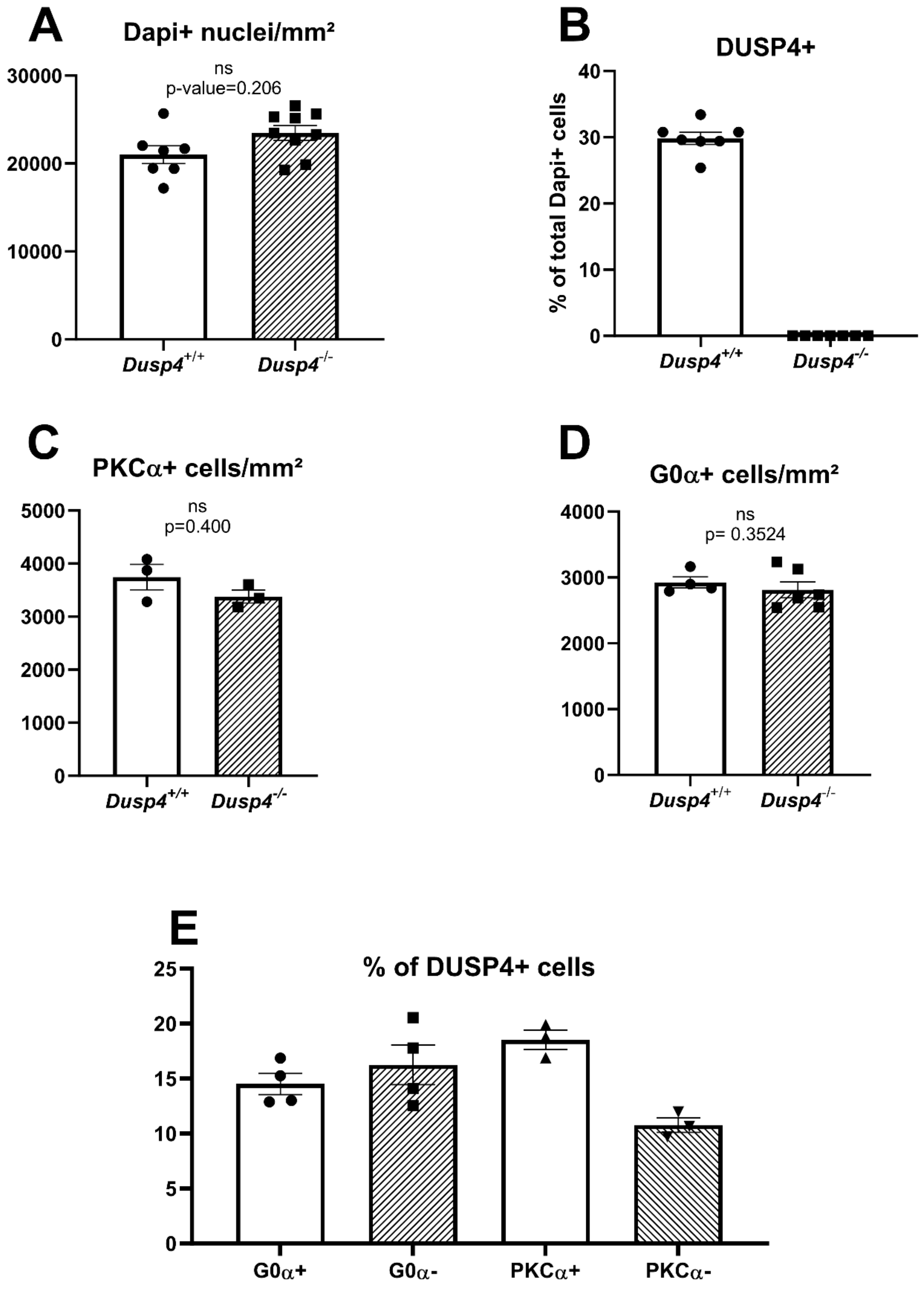


**Supplemental figure 2: Presence of DUSP4 in cells of adult mouse retina.** A: Quantification of Dapi positive nuclei in *Dusp4*$^{+/+}$ (n=7) and *Dusp4*$^{-/-}$ (n=9) retinas. B: Proportions of DUSP4 positive cells in *Dusp4*$^{+/+}$ (n=7) and *Dusp4*$^{-/-}$ (n=9) INLs. C: Quantification of PKCα+ cells in *Dusp4*$^{+/+}$ (n=3) and *Dusp4*$^{-/-}$ (n=3) INLs. D: Quantification of G0α+ cells in *Dusp4*$^{+/+}$ (n=4) and *Dusp4*$^{-/-}$ (n=6) INLs. E: Proportions of DUSP4+ cells co-staining (+) or not (-) with ON-BCs markers in *Dusp4*$^{+/+}$ retinas. Statistical significance between genotypes was tested using non parametric Mann-Whitney. ns: p-value>0.05.

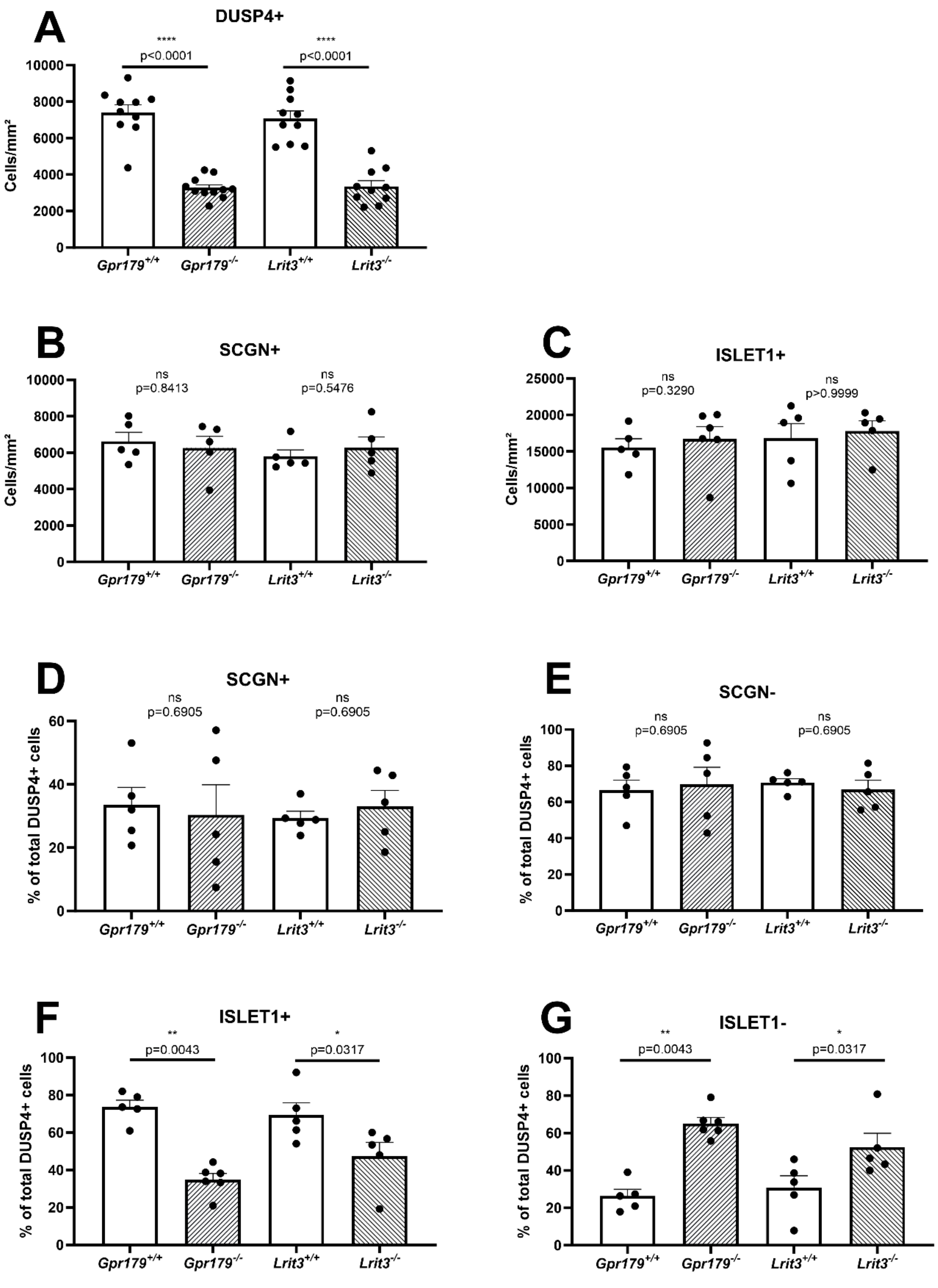


**Supplemental figure 3: ON-BCs specific loss of DUSP4 expression in cCSNB retinas.** A: Quantification of cells expressing DUSP4 in *Gpr179*$^{-/-}$ (n=11), *Lrit3*$^{-/-}$ (n=10) retinas and their wild-type littermates (n=10 each). B: Quantification of cells expressing SCGN in *Gpr179*$^{-/-}$ (n=5), *Lrit3*$^{-/-}$ (n=5) retinas and their wild-type littermates (n=5 each). C: Quantification of cells expressing ISLET1 in *Gpr179*$^{-}$

$^{/-}$ (n=6), *Lrit3*$^{-/-}$ (n=5) retinas and their wild-type littermates (n=5 each). D and E: Proportion of Dusp4 positive cells expressing (D) or not (E) SCGN in cCSNB and their wild-type littermates (n=5 each). F and G: Proportion of DUSP4 positive cells expressing (F) or not (G) ISLET1 in *Gpr179*$^{-/-}$ (n=6), *Lrit3*$^{-/-}$ (n=5) INLs and their wild-type littermates (n=5 each). Statistical significance between genotypes was tested using a non-parametric Mann-Whitney. ns: P-value>0.05; *: P-value≤0.05; **: p-value≤0.01; ****: p-value≤0.0001.

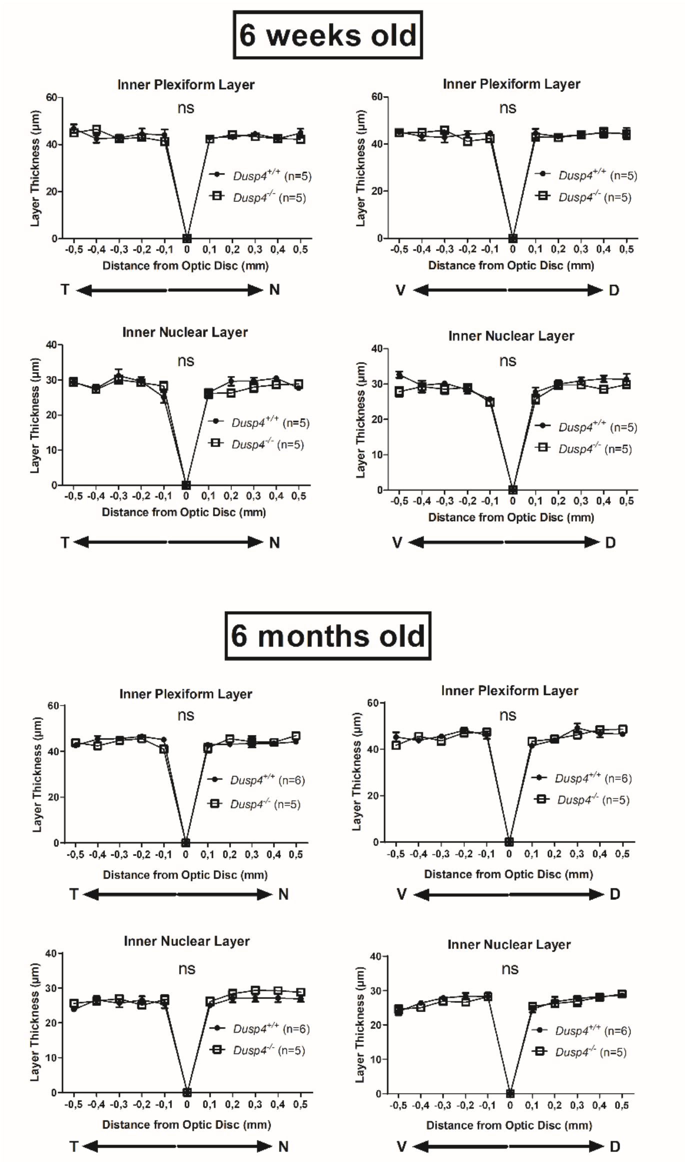


**Supplemental figure 4: No change of retinal layers thickness in *Dusp4*$^{-/-}$ mice.**

Inner plexiform layer and inner nuclear layer assessed using SD-OCT from 6-week-old and 6-month-old *Dusp4*$^{+/+}$ and *Dusp4*$^{-/-}$ mice. The layer thickness is represented according to the distance from the optic disc on both temporo-nasal (T⇔N) and ventro-dorsal (V⇔D) axis. Two-way ANOVA was used to test main effect of genotype. ns= non-significant, p-value>0.05.

| Gene | Relevant known pathway | Known retinal expression | Mouse model genetic alteration | MGI code | Retinal dopamin metabolism | Sensitivity to LIM | Key ERG phenotype | References |
|---|---|---|---|---|---|---|---|---|
| *Dusp4* | MAPK/ERK | ON-BCs, subset of OFF-BCs (subset of ACs?) | Constitutive Knock-out (Targeted homologous recombination) | MGI:2442191 | Reduced | Increased | Higher scotopic b-wave amplitude, lower scotopic OPs amplitude | Present study |
| *Pkcα* | MAPK/ERK | Rod-BCs | Constitutive Knock-out (Targeted null mutation) | MGI: 97595 | Reduced | Increased | Higher scotopic b-wave amplitude, lower scotopic OPs amplitude | 56, 54, present study |
| *Tpbg* | MAPK/ERK and Glutamate vesicle exocytosis | Rod-BCs | Constitutive Knock-out (Targeted null mutation) | MGI:5007391 | Reduced | Not investigated | Higher scotopic b-wave amplitude, lower scotopic OPs amplitude | 55, present study |
| *Gpr179* | Photoreceptors to ON-BCs signal transmission | ON-BCs | Constitutive Knock-out (Targeted stop codon insertion) | MGI:2443409 | Reduced | Increased | No scotopic b-wave, no scotopic OPs | 12, 34 |
| *Lrit3* | Photoreceptors to ON-BCs signal transmission | ON-BCs | Constitutive Knock-out (Targeted null mutation) | MGI:2685267 | Reduced | Increased | No scotopic b-wave, no scotopic OPs | 13, 33 |

**Supplemental table 1: Summary of the mouse lines used and the key findings of present study.** *Gpr179*$^{-/-}$ and *Lrit3*$^{-/-}$ mouse lines were used to assess the impact of ON-pathway defect upon *Dusp4* expression and localization. *Tpbg*$^{-/-}$ and *Pkcα*$^{-/-}$ mouse lines were used to support our hypothesis of reduced retinal levels of DA (i.e. potential myopia onset/sensitivity) despite a higher ON-BCs activity and helped us to provide clues about subsequent mechanisms.

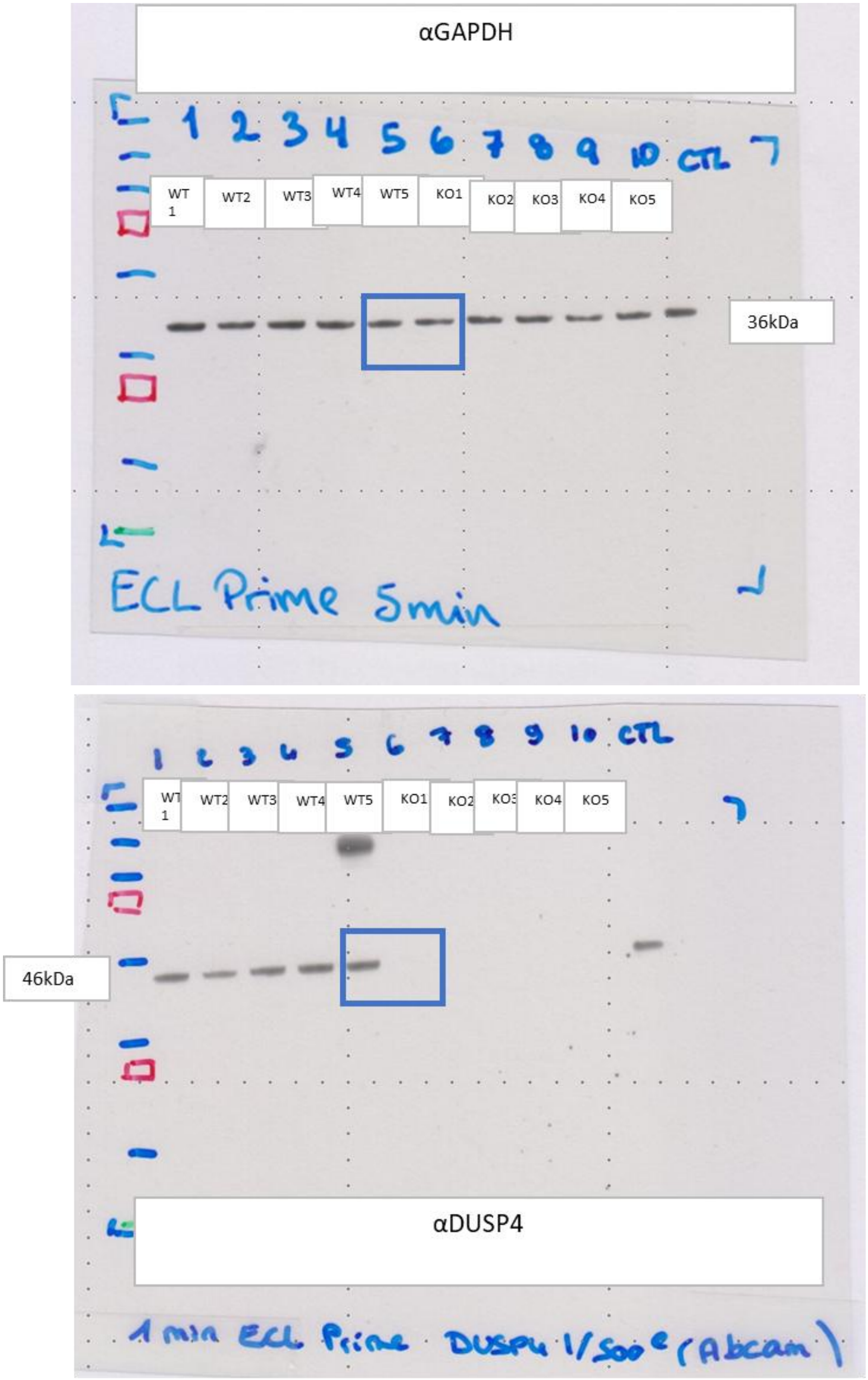


**Uncropped blots for Figure 2D.** WT= *Dusp4*$^{+/+}$, KO= *Dusp4*$^{-/-}$. Blue squares= Crops shown in original figures.

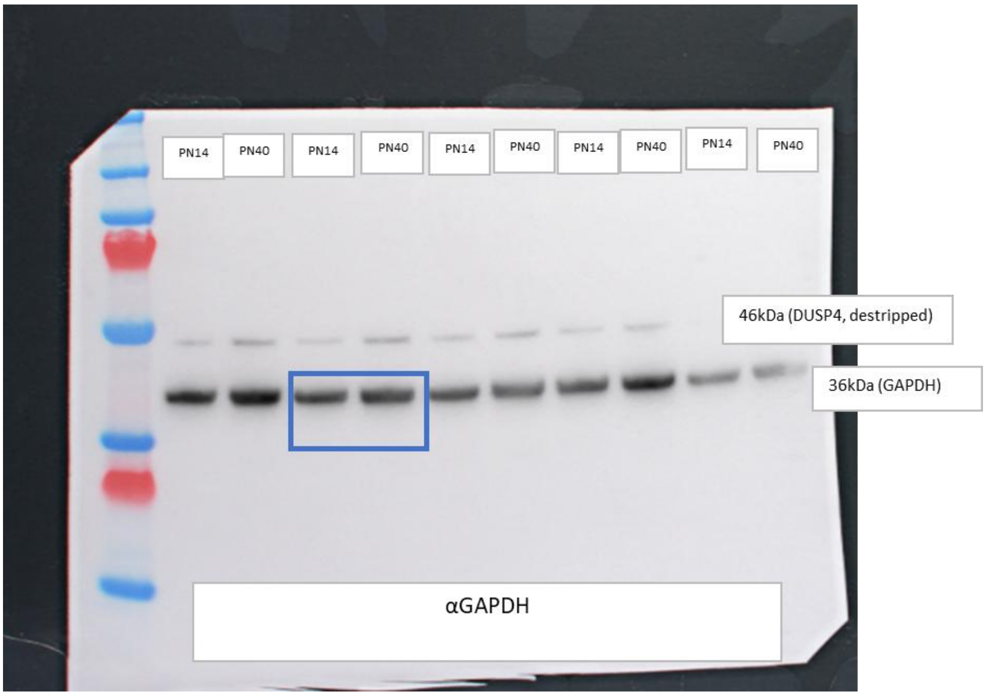


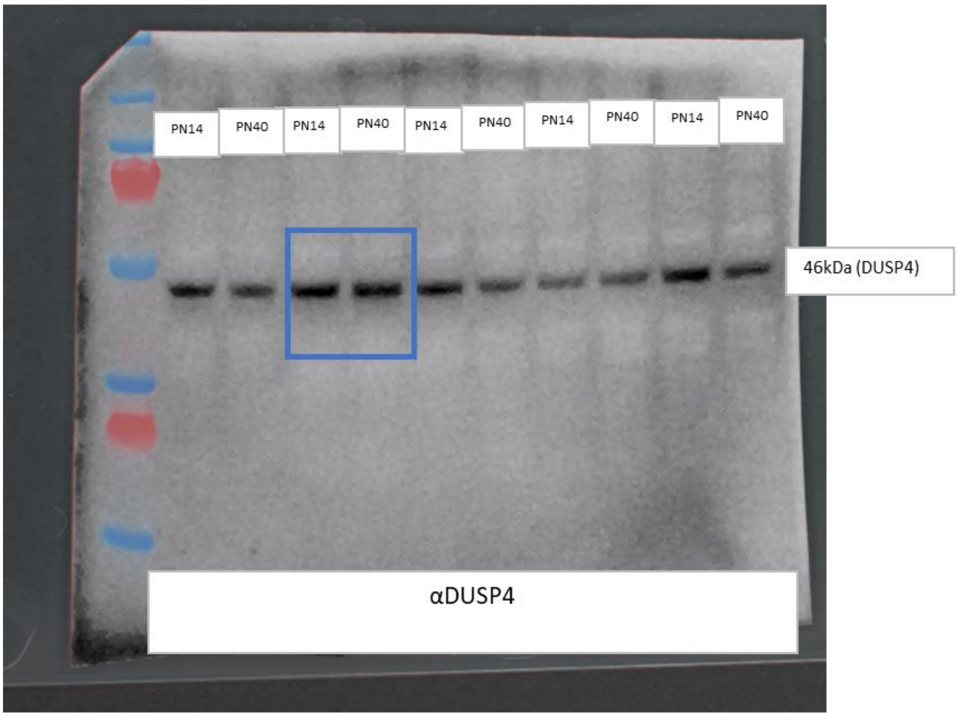


**Uncropped blots for Figure 2E.** PN14= *Dusp4*$^{+/+}$ retinas from post-natal day 14. PN40= *Dusp4*$^{+/+}$ retinas from post-natal day 40. Blue squares= Crops shown in original figures.

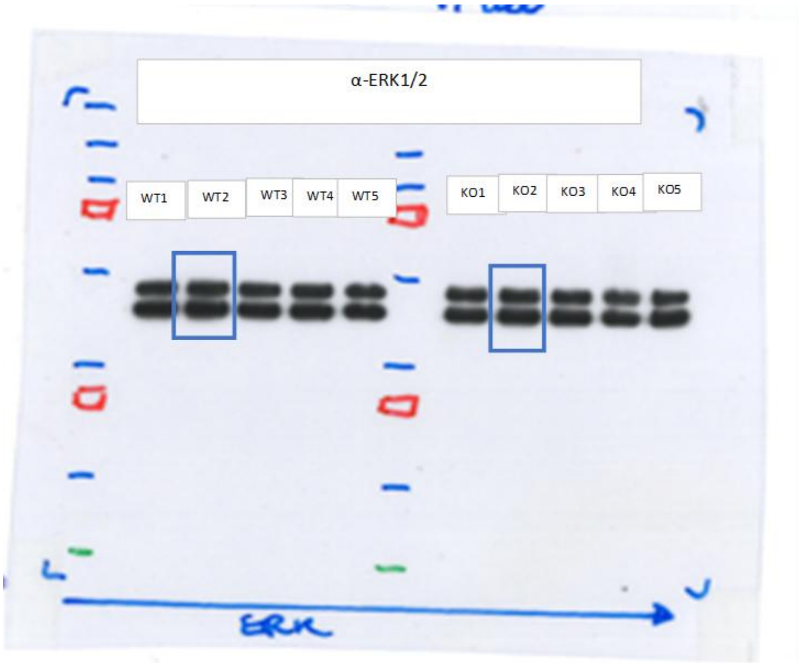


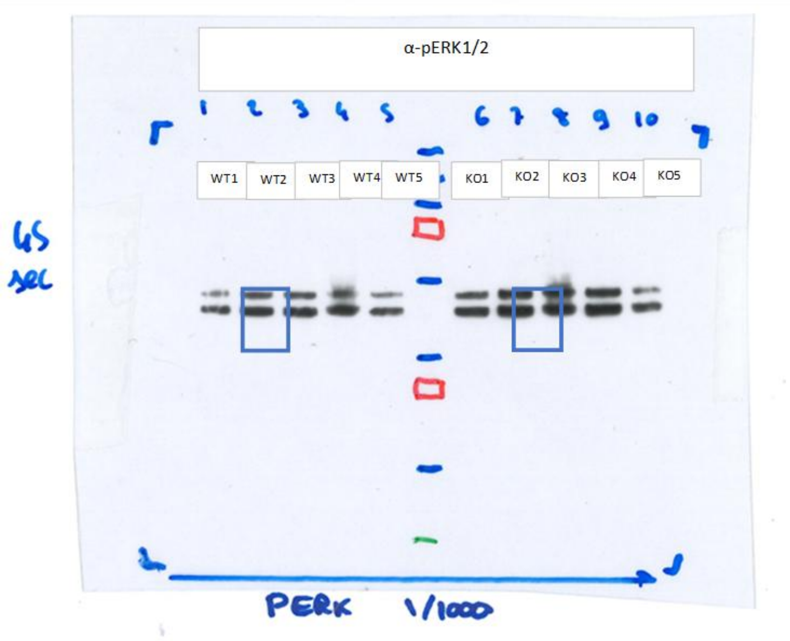


**Uncropped blots for Figure 2F.** WT= *Dusp4*$^{+/+}$, KO= *Dusp4*$^{-/-}$. Blue squares= Crops shown in original figures.

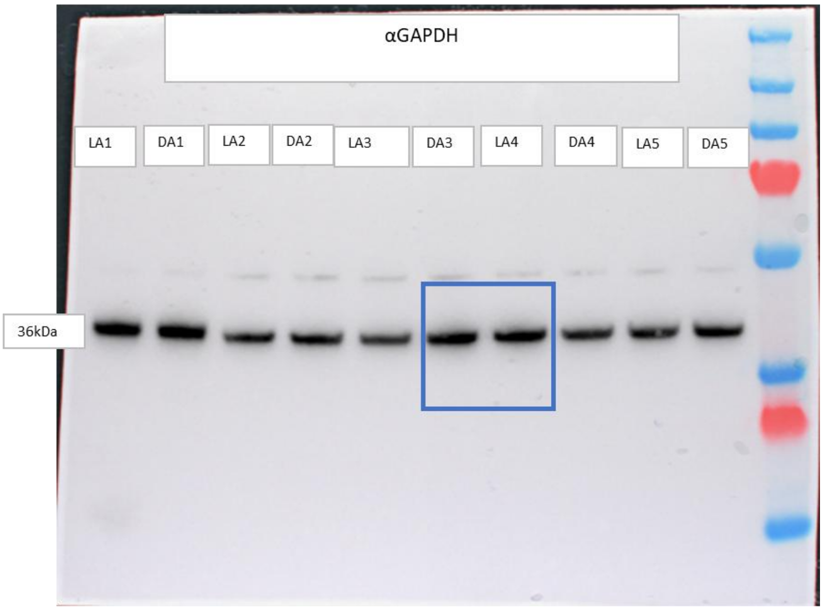


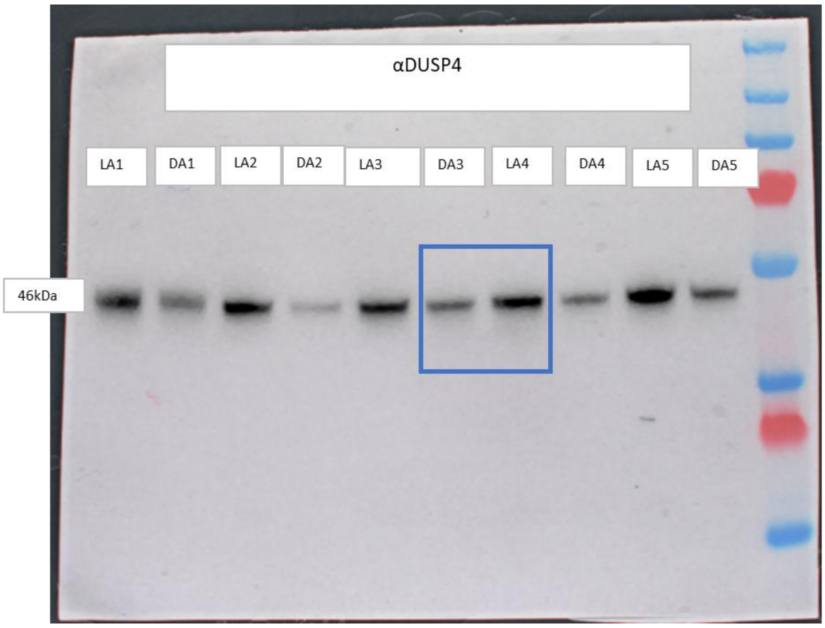


**Uncropped blots for Figure 2G.** LA= Light adapted *Dusp4*$^{+/+}$ retinas. DA= Dark adapted *Dusp4*$^{+/+}$ retinas. Blue squares= Crops shown in original figures.

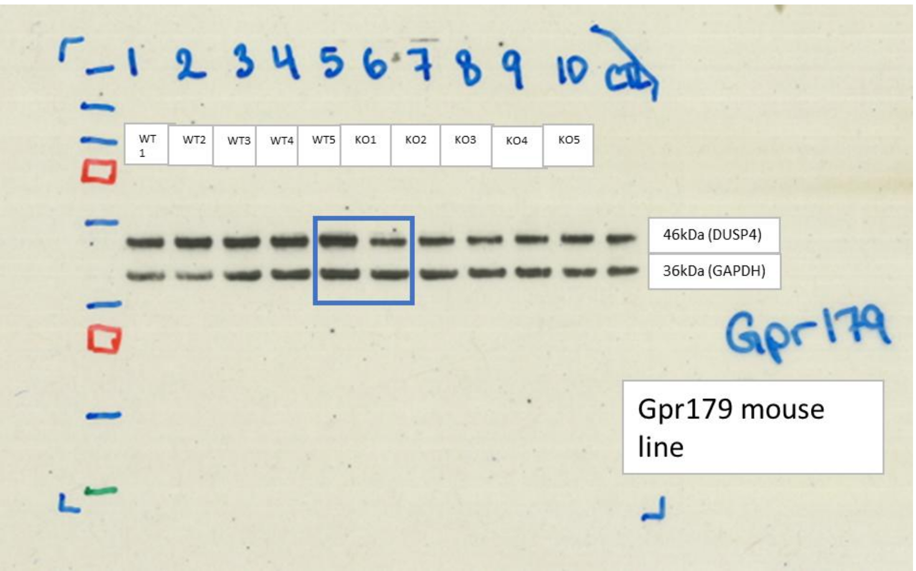


**Uncropped blots for Figure 3B.** WT= *Gpr179*$^{+/+}$, KO= *Gpr179*$^{-/-}$. Blue squares= Crops shown in original figures.

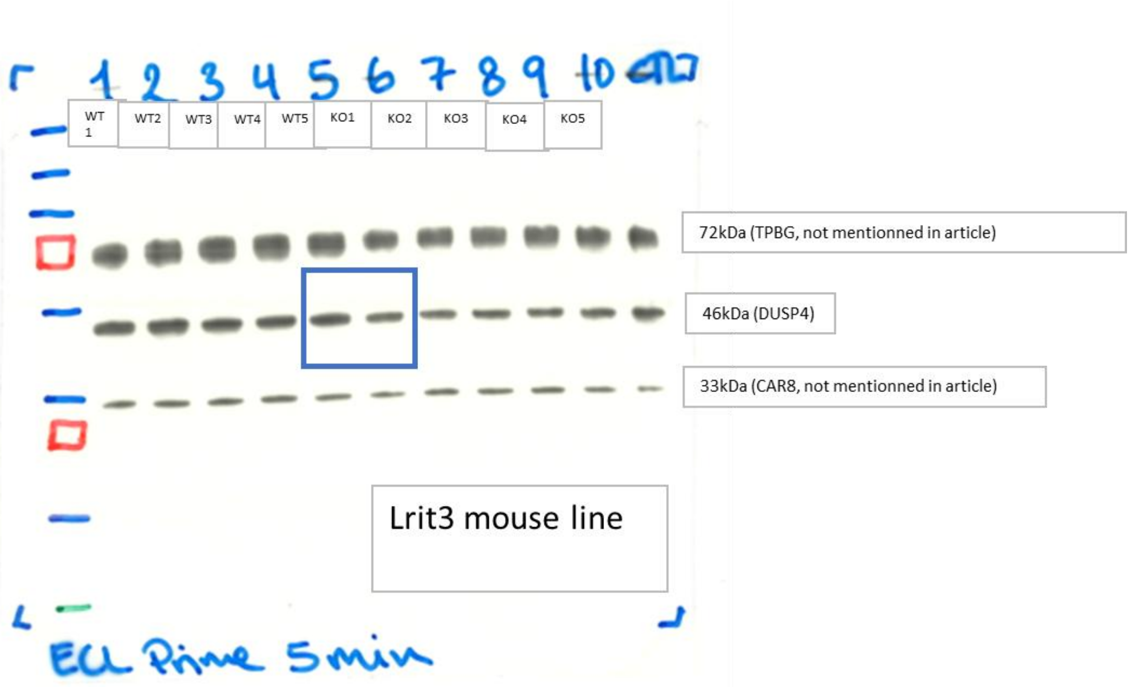


**Uncropped blots for Figure 3B.** WT= *Lrit3*$^{+/+}$, KO= *Lrit3*$^{-/-}$. Blue squares= Crops shown in original figures.

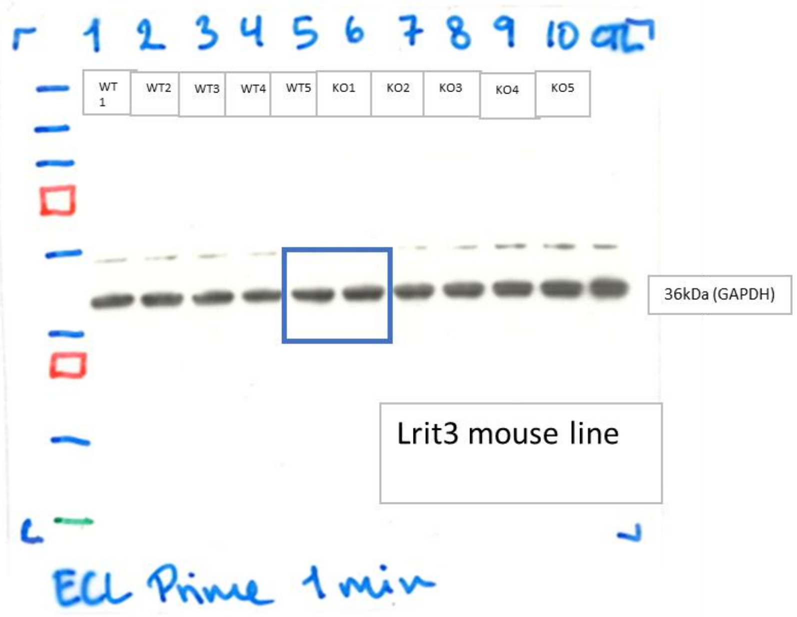


**Uncropped blots for Figure 3B.** WT= *Lrit3*$^{+/+}$, KO= *Lrit3*$^{-/-}$. Blue squares= Crops shown in original figures.
